# A prospective analysis of circulating plasma metabolomics and ovarian cancer risk

**DOI:** 10.1101/654962

**Authors:** Oana A. Zeleznik, A. Heather Eliassen, Peter Kraft, Elizabeth M. Poole, Bernard Rosner, Sarah Jeanfavre, Amy Deik, Kevin Bullock, Daniel Hitchcock, Julian Avila-Pancheco, Clary B. Clish, Shelley S. Tworoger

## Abstract

We assessed the association of pre-diagnostic plasma metabolites (N=420) with ovarian cancer risk. We included 252 cases and 252 matched controls from the Nurses’ Health Studies. Multivariable logistic regression was used to estimate odds ratios (OR) and 95% confidence intervals (CI) comparing the 90^th^-10^th^ percentile in metabolite levels, using permutation tests to account for testing multiple correlated hypotheses. Weighted gene co-expression network analysis (WGCNA) modules (n=10) and metabolite set enrichment analysis (MSEA; n=23) were also evaluated. Pseudouridine had the strongest statistical association with ovarian cancer risk overall (OR=2.56, 95%CI=1.48-4.45; p=0.001/adjusted-p=0.15). C36:2 phosphatidylcholine (PC) plasmalogen had the strongest statistical association with lower risk (OR=0.11, 95%CI=0.03-0.35; p<0.001/adjusted-p=0.06) and pseudouridine with higher risk (OR=9.84, 95%CI=2.89-37.82; p<0.001/adjusted-p=0.07) of non-serous tumors. Seven WGCNA modules and 15 classes were associated with risk at FDR≤0.20. Triacylglycerols (TAGs) showed heterogeneity by tumor aggressiveness (case-only heterogeneity-p<0.0001). TAG association with risk overall and serous tumors differed by acyl carbon content and saturation. Pseudouridine may be a novel risk factor for ovarian cancer. TAGs may also be important, particularly for rapidly fatal tumors, with associations differing by structural features. Validation in independent prospective studies and complementary experimental work to understand biological mechanisms is needed.

## Introduction

Ovarian cancer is the fifth leading cause of female cancer death in the U.S. (1). However, there are few known risk factors, such that current risk prediction models have a modest predictive capability. Thus, new strategies and research avenues to identify women at high risk are crucial to help prevent this highly fatal disease.

The metabolome consists of all metabolites, small molecules such as amino acids, carbohydrates and lipids, in a biological system (2), and reflects the integrated effect of genomics and environmental influences. Notably, advances in technology have led to precise measures of small molecule metabolites that are critical for growth and maintenance of cells in biologic fluids (3, 4). Several studies have identified specific metabolites as biomarkers of cancer risk. For example, branched chain amino acids were strongly associated with risk of pancreatic cancer (5) and lipid metabolites were strongly inversely associated with risk of aggressive prostate cancer (6). Further, prediagnostic serum concentrations of metabolites related to alcohol, vitamin E, and animal fats were modestly associated with ER^+^ breast cancer risk (7), while BMI-related metabolites were more strongly related to increased breast cancer risk (8). These findings support metabolomics profiling as a valuable strategy for identifying new markers of cancer risk.

Therefore, we used metabolomics assays to quantify several classes of circulating metabolites in plasma samples collected three to twenty-three years prior to ovarian cancer diagnosis within a nested case-control study, and, in an agnostic analysis, assessed their potential as biomarkers of ovarian cancer risk.

## Methods

### Study Population

We conducted nested case-control studies within the Nurses Health Studies (NHS (9) and NHSII (10)). The NHS was established in 1976 among 121,700 US female nurses aged 30–55 years, and NHSII was established in 1989 among 116,429 female nurses aged 25–42 years. Participants have been followed biennially by questionnaire to update information on exposure status and disease diagnoses. In 1989–1990, 32,826 NHS participants provided blood samples and completed a short questionnaire (9). Briefly, women arranged to have their blood drawn and shipped with an ice pack, via overnight courier, to our laboratory, where it was processed and separated into plasma, red blood cell, and white blood cell components and frozen in gasketed cryovials in the vapor phase of liquid nitrogen freezers. Between 1996 and 1999, 29,611 NHSII participants provided blood samples and completed a short questionnaire (10). Premenopausal women (*n*=18,521) who had not taken hormones, been pregnant, or lactated within the past 6 months provided blood samples drawn 7–9 days before the anticipated start of their next menstrual cycle (luteal phase). Other women (*n* = 11,090) provided a single 30-mL untimed blood sample. Samples were shipped and processed identically to the NHS samples.

Incident cases of epithelial ovarian cancer were identified through the biennial questionnaires or via linkage with the National Death Index. For women reporting a new ovarian cancer diagnosis or cases identified through death certificates, we obtained related medical records and pathology reports; for cases who had died we linked to the relevant cancer registry when medical records were unattainable. A gynecologic pathologist reviewed the records to confirm the diagnosis and abstract date of diagnosis, invasiveness, stage, and histotype (serous, poorly differentiated [PD], endometrioid, clear cell [CC], mucinous, other/unknown). Date of death was extracted from the death certificate. Participants diagnosed with invasive disease and who died within 3 years of diagnosis were defined as rapidly fatal cases (11).

Cases were diagnosed with ovarian cancer three years after blood collection until June 1, 2012 (NHS), or June 1, 2013 (NHSII). Two hundred fifty-three cases of invasive and borderline epithelial ovarian cancer (213 in NHS and 40 in NHSII) were confirmed by medical record review. Cases were matched to one control on: cohort (NHS, NHSII); menopausal status and hormone therapy use at blood draw (premenopausal, postmenopausal and hormone therapy use, postmenopausal and no hormone therapy use, missing/ unknown); menopausal status at diagnosis (premenopausal, postmenopausal, or unknown); age (±1 year), date of blood collection (±1 month); time of day of blood draw (±2 hours); and fasting status (>8 hours or ≤8 hours); women in NHSII who gave a luteal sample were matched on the luteal date (date of the next period minus date of blood draw, ±1 day).

The study protocol was approved by the institutional review boards of the Brigham and Women’s Hospital and Harvard T.H. Chan School of Public Health, and those of participating registries as required.

### Metabolite profiling

Plasma metabolites were profiled at the Broad Institute of MIT and Harvard (Cambridge, MA) using three complimentary liquid chromatography tandem mass spectrometry (LC-MS/MS) methods designed to measure polar metabolites and lipids as well as free fatty acids as described previously (12–15). For each method, pooled plasma reference samples were included every 20 samples and results were standardized using the ratio of the value of the sample to the value of the nearest pooled reference multiplied by the median of all reference values for the metabolite. Samples from the two cohorts were run together, with matched case-control pairs (as sets) distributed randomly within the batch, and the order of the case and controls within each pair randomly assigned. Therefore, the case and its control were always directly adjacent to each other in the analytic run, thereby limiting variability in platform performance across matched case-control pairs. In addition, 64 quality control (QC) samples, to which the laboratory was blinded, were also profiled. These were randomly distributed among the participants’ samples.

Hydrophilic interaction liquid chromatography (HILIC) analyses of water soluble metabolites in the positive ionization mode were conducted using an LC-MS system comprised of a Shimadzu Nexera X2 U-HPLC (Shimadzu Corp.; Marlborough, MA) coupled to a Q Exactive mass spectrometer (Thermo Fisher Scientific; Waltham, MA). Metabolites were extracted from plasma (10 μL) using 90 μL of acetonitrile/methanol/formic acid (74.9:24.9:0.2 v/v/v) containing stable isotope-labeled internal standards (valine-d8, Sigma-Aldrich; St. Louis, MO; and phenylalanine-d8, Cambridge Isotope Laboratories; Andover, MA). The samples were centrifuged (10 min, 9,000 × g, 4°C), and the supernatants were injected directly onto a 150 × 2 mm, 3 μm Atlantis HILIC column (Waters; Milford, MA). The column was eluted isocratically at a flow rate of 250 μL/min with 5% mobile phase A (10 mM ammonium formate and 0.1% formic acid in water) for 0.5 minute followed by a linear gradient to 40% mobile phase B (acetonitrile with 0.1% formic acid) over 10 minutes. MS analyses were carried out using electrospray ionization in the positive ion mode using full scan analysis over 70-800 m/z at 70,000 resolution and 3 Hz data acquisition rate. Other MS settings were: sheath gas 40, sweep gas 2, spray voltage 3.5 kV, capillary temperature 350°C, S-lens RF 40, heater temperature 300°C, microscans 1, automatic gain control target 1e6, and maximum ion time 250 ms.

Plasma lipids were profiled using a Shimadzu Nexera X2 U-HPLC (Shimadzu Corp.; Marlborough, MA). Lipids were extracted from plasma (10 μL) using 190 μL of isopropanol containing 1,2-didodecanoyl-sn-glycero-3-phosphocholine (Avanti Polar Lipids; Alabaster, AL). After centrifugation, supernatants were injected directly onto a 100 × 2.1 mm, 1.7 μm ACQUITY BEH C8 column (Waters; Milford, MA). The column was eluted isocratically with 80% mobile phase A (95:5:0.1 vol/vol/vol 10mM ammonium acetate/methanol/formic acid) for 1 minute followed by a linear gradient to 80% mobile-phase B (99.9:0.1 vol/vol methanol/formic acid) over 2 minutes, a linear gradient to 100% mobile phase B over 7 minutes, then 3 minutes at 100% mobile-phase B. MS analyses were carried out using electrospray ionization in the positive ion mode using full scan analysis over 200–1100 m/z at 70,000 resolution and 3 Hz data acquisition rate. Other MS settings were: sheath gas 50, in source CID 5 eV, sweep gas 5, spray voltage 3 kV, capillary temperature 300°C, S-lens RF 60, heater temperature 300°C, microscans 1, automatic gain control target 1e6, and maximum ion time 100 ms. Lipid identities were denoted by total acyl carbon number and total double bond number.

Metabolites of intermediate polarity, including free fatty acids and bile acids, were profiled using a Nexera X2 U-HPLC (Shimadzu Corp.; Marlborough, MA) coupled to a Q Exactive (Thermo Fisher Scientific; Waltham, MA). Plasma samples (30 μL) were extracted using 90 μL of methanol containing PGE2-d4 as an internal standard (Cayman Chemical Co.; Ann Arbor, MI) and centrifuged (10 min, 9,000 × g, 4°C). The supernatants (10 μL) were injected onto a 150 × 2.1 mm ACQUITY BEH C18 column (Waters; Milford, MA). The column was eluted isocratically at a flow rate of 450 μL/min with 20% mobile phase A (0.01% formic acid in water) for 3 minutes followed by a linear gradient to 100% mobile phase B (0.01% acetic acid in acetonitril) over 12 minutes. MS analyses were carried out using electrospray ionization in the negative ion mode using full scan analysis over m/z 70-850. Additional MS settings are: ion spray voltage, −3.5 kV; capillary temperature, 320°C; probe heater temperature, 300 °C; sheath gas, 45; auxiliary gas, 10; and S-lens RF level 60.

Raw data from orbitrap mass spectrometers were processed using TraceFinder 3.3 software (Thermo Fisher Scientific; Waltham, MA) and Progenesis QI (Nonlinear Dynamics; Newcastle upon Tyne, UK) and targeted data from the QTRAP 5500 system were processed using MultiQuant (version 2.1, SCIEX; Framingham, MA). For each method, metabolite identities were confirmed using authentic reference standards or reference samples.

In total, 608 known metabolites were measured in this study. Metabolites with a coefficient of variation (CV) among blinded QC samples higher than 25%, or an intraclass correlation coefficient (ICC) <0.4 were excluded from this analysis (N=132, **Supplementary Table 1**). Furthermore, metabolites not passing our previously conducted processing delay pilot study (15) were excluded from this analysis (N=56, **Supplementary Table 1**). All metabolites (N=420, **Supplementary Table 1**) included in the analysis exhibited good reproducibility within person over one year (15). 197 metabolites had no missing values among participant samples. Missing values in metabolites (N=211) with less than 10% missingness were imputed with 1/2 of the minimum value measured for that metabolite. We included a missing value indicator for metabolites (N=12) with more than 10% missingness (see statistical analysis section for further details).

After these quality control exclusions total of 420 metabolites including amino acids, amino acids derivatives, amines, lipids, fatty acids, and bile acids were analyzed in this study. Continuous metabolite values were transformed to probit scores for all analyses to reduce the influence of skewed distributions and heavy tails on the results and to scale the measured metabolite values to the same range.

### Statistical analysis

#### Identification of individual metabolites associated with risk

Conditional logistic regression was used to evaluate metabolite associations, modeled continuously, with risk of overall ovarian cancer. For metabolites with more than 10% missing values, we added a missing indicator term to the regression model. We present the odds ratios (OR) and 95% confidence intervals (95% CI) for an increase from the 10^th^ to 90^th^ percentile in metabolite levels or the indicator variable as appropriate.

In a sensitivity analysis, we compared conditional logistic regression to unconditional logistic regression adjusting for the matching factors and found similar results (data not shown). Thus, subsequent analyses by histotype, rapidly fatal status and time between blood collection and diagnosis were conducted using unconditional logistic regression adjusting for the matching factors, allowing the use of all controls.

We conducted stratified analyses restricting separately to serous/PD tumors (cases=176/controls=252), endometrioid/CC tumors (cases=34/controls=252), premenopausal (cases=82/controls=82) and postmenopausal women (cases=137/controls=137) at blood collection, to participants diagnosed 3-11 years (cases=121/controls=252) and 12-23 years after blood collection (cases=131/controls=252), to rapidly fatal cases (defined as death occurring within 3 years of diagnosis; cases=86/controls=252), and to less aggressive tumors (defined as death occurring at least 3 years after diagnosis; cases=138/controls=252). All models were adjusted for matching factors and established ovarian cancer risk factors: duration of oral contraceptive use (none or <3 months, 3 months to 3 years, 3 years to 5 years, more than 5 years), tubal ligation (yes/no) and parity (no children, 1 child, 2 children, 3 children, 4+ children). We calculated heterogeneity by time to diagnosis and tumor aggressiveness using case-only analyses and by menopausal status at blood collection by introducing an interaction term between the metabolite and menopausal status.

A permutation test (N=5000) was used to control the family-wise error rate (i.e. account for multiple testing) while accounting for the correlation structure of metabolites. Case-control status was permuted within a matched case-control pair for conditional logistic regression analyses. To account for all controls included in the subtype analyses using unconditional logistic regression, each control was matched to a case within that analysis, while preserving the initial matching criteria as much as possible. The smallest p-value across all tested metabolites in each permutation run was recorded. The permutation p-value for test of the overall null (no metabolite is associated with ovarian cancer) was estimated as k/(5,001), where k is the number of permutations where the smallest p-value (across all metabolites) was smaller than the smallest observed p-value. We estimated the permutation adjusted p-value for each metabolite by using the stepdown min P approach by Westfall and Young (16) implemented in the R package ***NPC*** which is based on the previously computed permutation p-values and accounts for the correlation structure among metabolites.

#### Identification of groups of metabolites associated with risk

Metabolite Set Enrichment Analysis (MSEA) (17), implemented in the R package ***FGSEA*** (18), was used to identify groups of molecularly or biologically similar metabolites that were enriched among the metabolites associated with risk of overall ovarian cancer and histotypes. This method ranks the metabolites by the estimated beta coefficient of the association with risk and uses this metric to identify enriched metabolite groups at the two extremes of the distribution of beta estimates (positive/inverse associations). Weighted Gene Co-expression Network Analysis (WGCNA) (19), implemented in the R package ***WGCNA*** uses hierarchical clustering to identify groups of correlated metabolites, called metabolite modules, which reflect a scale-free network topology of the measured metabolites (20). Modules were derived based on control samples only. Each module was summarized by its first principal component (PC) among all analyzed samples. A score was derived for each metabolite module based on the linear combination of measured metabolite values weighted by their corresponding loadings on the first PC summarizing the module. The score was subsequently used in conditional/unconditional logistic regressions to assess associations with risk of ovarian cancer overall and by histotypes. We report nominal p-values and false discovery rates (FDR) (21) for all metabolite groups and metabolite modules. All analyses were performed using the statistical computing language R (22).

## Results

### Study population

Of the 252 cases in the analysis, 176 cases were diagnosed with serous/PD tumors while 34 were classified as endometrioid/CC tumors (**Table 1**). The remaining cases were of mucinous or other types. Mean follow-up time was 12.3 years. Of the 252 cases, 86 represented rapidly fatal tumors with death within 3 years of diagnosis. Distributions of ovarian cancer risk factors were generally in the expected directions for cases and controls.

**Table 1:** Characteristics of overall, serous/poorly differentiated (PD) and endometrioid/clear cell (CC) ovarian cancer (OC) cases, rapidly fatal tumors, and all controls at time of blood collection.

|  | All Controls<br>(N = 252) | Overall OC<br>(N = 252) | Serous/PD OC<br>(N = 176) | Endometrioid/CC OC<br>(N = 34) | Other histotypes<br>(N = 42) | Rapidly fatal<br>tumors (N=86) |
| --- | --- | --- | --- | --- | --- | --- |
| **Mean (SD)** |  |  |  |  |  |  |
| Age at blood draw* | 55.6 (7.8) | 55.5 (7.9) | 55.3 (7.9) | 54.0 (8.1) | 57.8 (7.5) | 58.5 (6.8) |
| Time to diagnosis (years) | - | 12.3 (5.2) | 12.8 (5.3) | 12.1 (5.1) | 10.9 (4.7) | 12.7 (5.2) |
| Age at diagnosis (years) | - | 69.7 (9.7) | 68.1 (9.8) | 66.0 (9.8) | 68.7 (9.2) | 71.2 (8.7) |
| **N (Percent)** |  |  |  |  |  |  |
| Tumor morphology |  |  |  |  |  |  |
| Invasive | - | 227 (90) | 163 (93) | 33 (97) | 31 (74) | 86 (100) |
| Borderline | - | 22 (9) | 13 (7) | 1 (3) | 8 (19) | 0 (0) |
| Unknown | - | 3 (1) | 0 (0) | 0 (0) | 3 (7) | 0 (0) |
| Menopausal status blood draw* |  |  |  |  |  |  |
| Premenopausal | 82 (33) | 82 (33) | 56 (32) | 16 (47) | 10 (24) | 14 (16) |
| Postmenopausal, No HT use | 71 (28) | 68 (27) | 47 (27) | 5 (15) | 16 (38) | 29 (34) |
| Postmenopausal, HT use | 66 (26) | 69 (27) | 48 (27) | 8 (24) | 13 (31) | 34 (40) |
| Unknown | 33 (13) | 33 (13) | 25 (14) | 5 (15) | 3 (7) | 9 (10) |
| Cohort* |  |  |  |  |  |  |
| NHS | 212 (84) | 212 (84) | 147 (84) | 27 (79) | 38 (90) | 79 (92) |
| NHS II | 40 (16) | 40 (16) | 29 (16) | 7 (21) | 4 (10) | 7 (8) |
| Oral contraceptive use duration |  |  |  |  |  |  |
| None or <3 months | 123 (49) | 118 (47) | 81 (46) | 18 (53) | 19 (45) | 48 (56) |
| 3 months to 3 years | 33 (13) | 32 (13) | 22 (12) | 3 (9) | 7 (17) | 10 (12) |
| 3 to 5 years | 45 (18) | 63 (25) | 46 (26) | 8 (24) | 9 (21) | 16 (19) |
| 5+ years | 51 (20) | 39 (15) | 27 (15) | 5 (15) | 7 (17) | 12 (14) |
| Parity |  |  |  |  |  |  |
| No children | 12 (5) | 24 (10) | 16 (9) | 5 (15) | 3 (7) | 7 (8) |
| 1 child | 11 (4) | 13 (5) | 8 (5) | 1 (3) | 4 (10) | 2 (2) |
| 2 children | 72 (29) | 89 (35) | 60 (34) | 15 (44) | 14 (33) | 23 (27) |
| 3 children | 77 (31) | 65 (26) | 45 (26) | 9 (26) | 11 (26) | 25 (29) |
| 4+ children | 80 (32) | 61 (24) | 47 (27) | 4 (12) | 10 (24) | 29 (34) |
| Tubal ligation |  |  |  |  |  |  |
| Yes | 43 (17) | 39 (15) | 30 (17) | 5 (15) | 34 (10) | 17 (20) |
| * matching factors; HT hormone therapy |  |  |  |  |  |  |

### Measured metabolites and their association with ovarian cancer risk

Metabolite profiling resulted in 608 measured metabolites with 420 (69%) metabolites passing our QC filtering criteria, which included: 159 lipids, 158 amino acids, amino acids derivatives amines and cationic metabolites, and 103 free fatty acids, bile acids and lipid mediators. Eight metabolites were associated with risk of overall ovarian cancer at a nominal p-value ≤0.01 (**Table 2A**, **Figure 1** and **Supplementary Table 1**). Odds ratios for an increase from the 10^th^ to the 90^th^ percentile of metabolites levels for these metabolites ranged between 0.49 and 2.56. The top three metabolites associated with risk were pseudouridine (OR=2.56, 95% CI=1.48-4.45; p-value=0.001), C18:0 sphingomyelin (SM) (OR=2.1, 95% CI=1.26-3.49; p-value=0.004) and 4-acetamidobutanoate (OR=2.1, 95% CI=1.24-3.56; p-value=0.006). Pseudouridine had an adjusted p=0.15 while all other metabolites had adjusted p>0.5. The test of the global null hypothesis that no metabolite was associated with risk had p=0.15.

**Table 2:**
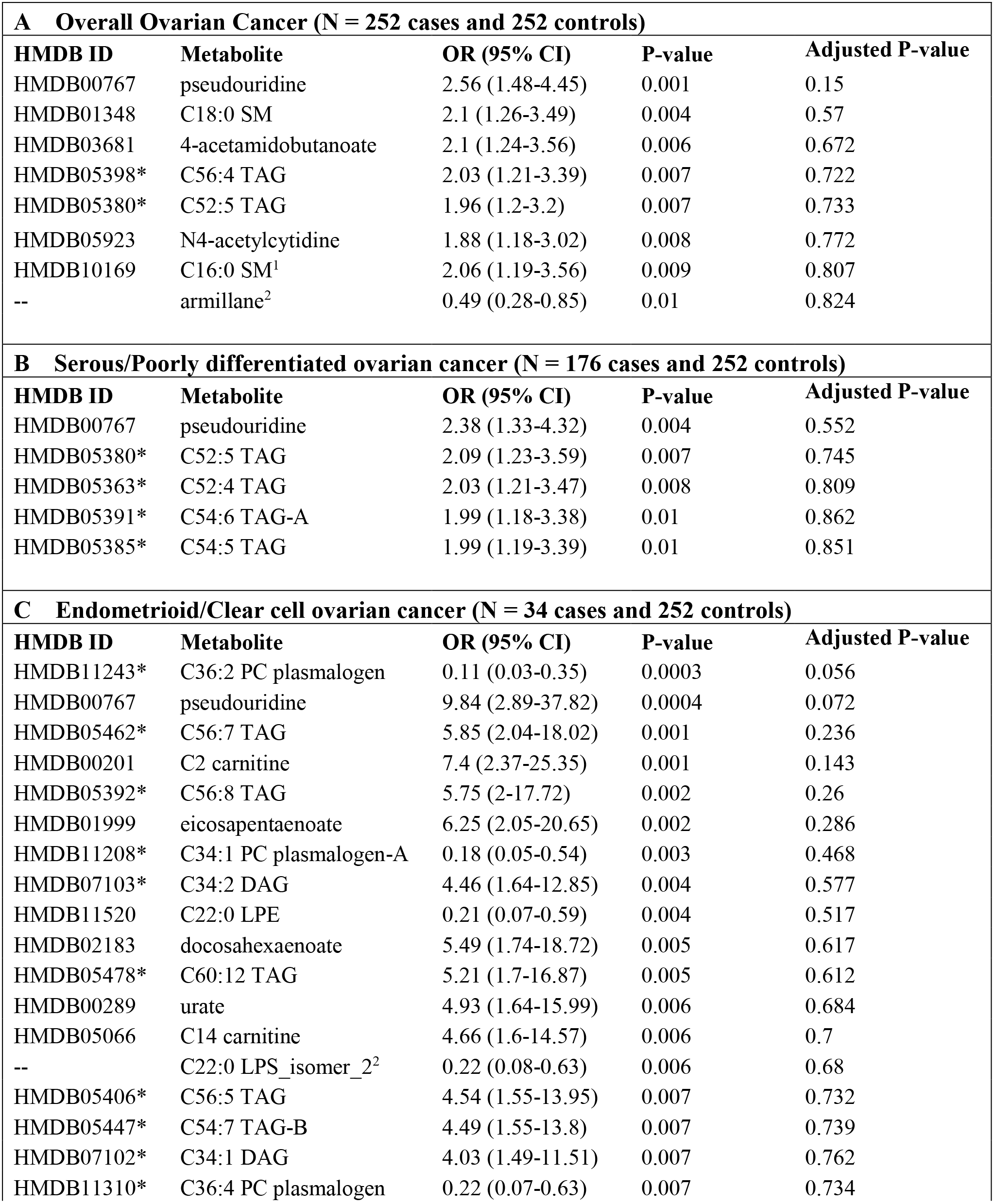
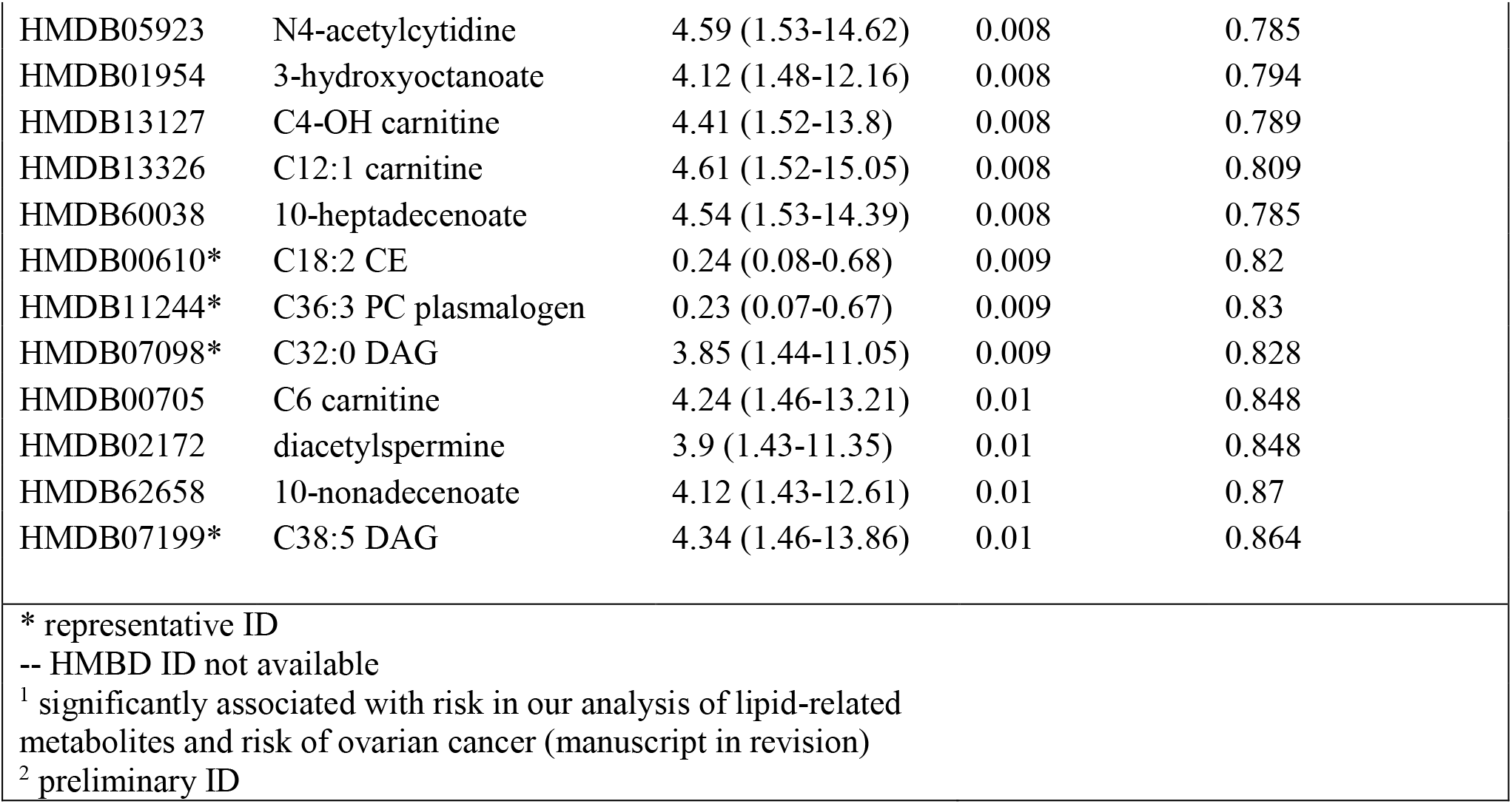
Odds ratio (OR) for an increase from the 10^th^ to the 90^th^ percentile of metabolite levels and 95% confidence intervals (CI) of associations with risk of overall, serous/poorly differentiated and endometrioid/clear cell ovarian cancer. Results with p-values ≤0.01 are shown.

| **A Overall Ovarian Cancer (N = 252 cases and 252 controls)** |  |  |  |  |
| --- | --- | --- | --- | --- |
| **HMDB ID** | **Metabolite** | **OR (95% CI)** | **P-value** | **Adjusted P-value** |
| HMDB00767 | pseudouridine | 2.56 (1.48-4.45) | 0.001 | 0.15 |
| HMDB01348 | C18:0 SM | 2.1 (1.26-3.49) | 0.004 | 0.57 |
| HMDB03681 | 4-acetamidobutanoate | 2.1 (1.24-3.56) | 0.006 | 0.672 |
| HMDB05398* | C56:4 TAG | 2.03 (1.21-3.39) | 0.007 | 0.722 |
| HMDB05380* | C52:5 TAG | 1.96 (1.2-3.2) | 0.007 | 0.733 |
| HMDB05923 | N4-acetylcytidine | 1.88 (1.18-3.02) | 0.008 | 0.772 |
| HMDB10169 | C16:0 SM<sup>1</sup> | 2.06 (1.19-3.56) | 0.009 | 0.807 |
| -- | armillane<sup>2</sup> | 0.49 (0.28-0.85) | 0.01 | 0.824 |
| **B Serous/Poorly differentiated ovarian cancer (N = 176 cases and 252 controls)** |  |  |  |  |
| **HMDB ID** | **Metabolite** | **OR (95% CI)** | **P-value** | **Adjusted P-value** |
| HMDB00767 | pseudouridine | 2.38 (1.33-4.32) | 0.004 | 0.552 |
| HMDB05380* | C52:5 TAG | 2.09 (1.23-3.59) | 0.007 | 0.745 |
| HMDB05363* | C52:4 TAG | 2.03 (1.21-3.47) | 0.008 | 0.809 |
| HMDB05391* | C54:6 TAG-A | 1.99 (1.18-3.38) | 0.01 | 0.862 |
| HMDB05385* | C54:5 TAG | 1.99 (1.19-3.39) | 0.01 | 0.851 |
| **C Endometrioid/Clear cell ovarian cancer (N = 34 cases and 252 controls)** |  |  |  |  |
| **HMDB ID** | **Metabolite** | **OR (95% CI)** | **P-value** | **Adjusted P-value** |
| HMDB11243* | C36:2 PC plasmalogen | 0.11 (0.03-0.35) | 0.0003 | 0.056 |
| HMDB00767 | pseudouridine | 9.84 (2.89-37.82) | 0.0004 | 0.072 |
| HMDB05462* | C56:7 TAG | 5.85 (2.04-18.02) | 0.001 | 0.236 |
| HMDB00201 | C2 carnitine | 7.4 (2.37-25.35) | 0.001 | 0.143 |
| HMDB05392* | C56:8 TAG | 5.75 (2-17.72) | 0.002 | 0.26 |
| HMDB01999 | eicosapentaenoate | 6.25 (2.05-20.65) | 0.002 | 0.286 |
| HMDB11208* | C34:1 PC plasmalogen-A | 0.18 (0.05-0.54) | 0.003 | 0.468 |
| HMDB07103* | C34:2 DAG | 4.46 (1.64-12.85) | 0.004 | 0.577 |
| HMDB11520 | C22:0 LPE | 0.21 (0.07-0.59) | 0.004 | 0.517 |
| HMDB02183 | docosahexaenoate | 5.49 (1.74-18.72) | 0.005 | 0.617 |
| HMDB05478* | C60:12 TAG | 5.21 (1.7-16.87) | 0.005 | 0.612 |
| HMDB00289 | urate | 4.93 (1.64-15.99) | 0.006 | 0.684 |
| HMDB05066 | C14 carnitine | 4.66 (1.6-14.57) | 0.006 | 0.7 |
| -- | C22:0 LPS_isomer_2<sup>2</sup> | 0.22 (0.08-0.63) | 0.006 | 0.68 |
| HMDB05406* | C56:5 TAG | 4.54 (1.55-13.95) | 0.007 | 0.732 |
| HMDB05447* | C54:7 TAG-B | 4.49 (1.55-13.8) | 0.007 | 0.739 |
| HMDB07102* | C34:1 DAG | 4.03 (1.49-11.51) | 0.007 | 0.762 |
| HMDB11310* | C36:4 PC plasmalogen | 0.22 (0.07-0.63) | 0.007 | 0.734 |
| HMDB05923 | N4-acetylcytidine | 4.59 (1.53-14.62) | 0.008 | 0.785 |
| HMDB01954 | 3-hydroxyoctanoate | 4.12 (1.48-12.16) | 0.008 | 0.794 |
| HMDB13127 | C4-OH carnitine | 4.41 (1.52-13.8) | 0.008 | 0.789 |
| HMDB13326 | C12:1 carnitine | 4.61 (1.52-15.05) | 0.008 | 0.809 |
| HMDB60038 | 10-heptadecenoate | 4.54 (1.53-14.39) | 0.008 | 0.785 |
| HMDB00610* | C18:2 CE | 0.24 (0.08-0.68) | 0.009 | 0.82 |
| HMDB11244* | C36:3 PC plasmalogen | 0.23 (0.07-0.67) | 0.009 | 0.83 |
| HMDB07098* | C32:0 DAG | 3.85 (1.44-11.05) | 0.009 | 0.828 |
| HMDB00705 | C6 carnitine | 4.24 (1.46-13.21) | 0.01 | 0.848 |
| HMDB02172 | diacetylspermine | 3.9 (1.43-11.35) | 0.01 | 0.848 |
| HMDB62658 | 10-nonadecenoate | 4.12 (1.43-12.61) | 0.01 | 0.87 |
| HMDB07199* | C38:5 DAG | 4.34 (1.46-13.86) | 0.01 | 0.864 |
| * representative ID<br>-- HMBD ID not available<br><sup>1</sup> significantly associated with risk in our analysis of lipid-related metabolites and risk of ovarian cancer (manuscript in revision)<br><sup>2</sup> preliminary ID |  |  |  |  |

**Figure 1:**
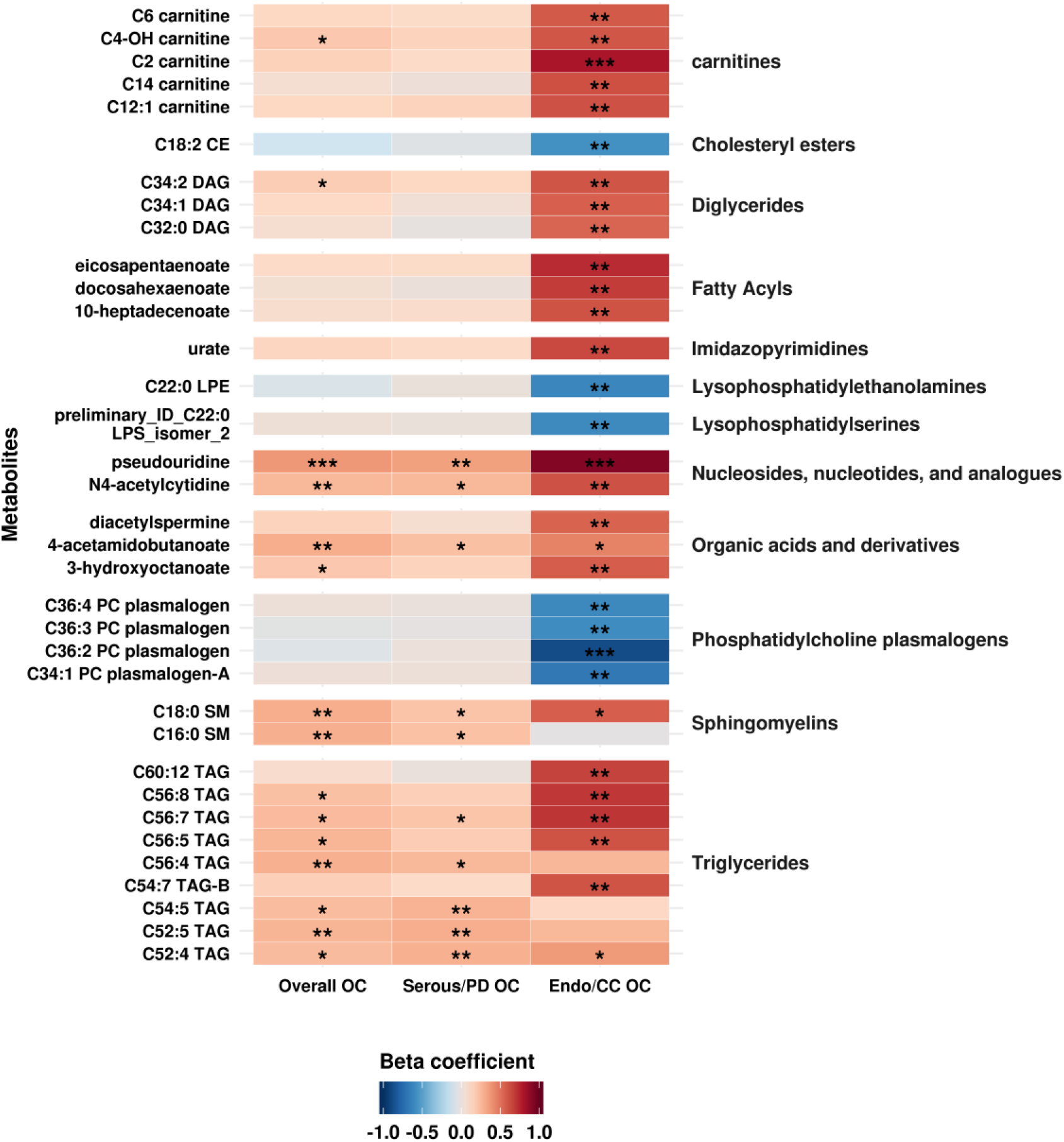
Beta coefficients of the association between metabolites and overall OC, serous/poorly differentiated OC (Serous/PD OC) and endometrioid/clear cell OC (Endo/CC OC). Coefficients with a p-value ≤0.01 in any of the analyses are shown. Shades of red represent positive coefficients while shades of blue indicate negative coefficients. Significance of the association is overlaid on the heat map and marked as follows: * p-values≤0.1, ** p-values≤0.01, *** p-values≤0.001; all other p-values are >0.1

Five metabolites were associated with risk of serous/PD tumors at a nominal p-value ≤0.01 (**Table 2B**, **Figure 1** and **Supplementary Table 2**). Odds ratios for an increase from the 10^th^ to the 90^th^ percentile of metabolites levels for these metabolites ranged between 1.99 and 2.38. The top three metabolites were pseudouridine (OR=2.38, 95% CI=1.33-4.32; p-value=0.004), C52:5 triglyceride (TAG) (OR=2.09, 95% CI=1.23-3.59; p-value=0.007) and C52:4 TAG (OR=2.03, 95% CI=1.21-3.47; p=0.008). However, none of the metabolites remained significant after accounting for multiple comparisons via permutation (adjusted p-value >0.55). The test of the global null hypothesis that no metabolite was associated with risk had p=0.55.

Thirty metabolites were associated with risk of endometrioid/CC tumors at a nominal p-value ≤0.01 (**Table 2C**, **Figure 1** and **Supplementary Table 2**). Odds ratios for an increase from 10^th^ to the 90^th^ percentile of metabolites levels for these metabolites ranged between 0.11 and 0.24 for inverse associations, and between 3.85 and 9.84 for positive associations. The top three metabolites positively associated with risk were pseudouridine (OR=9.84, 95% CI=2.89-37.82; p=0.0003), C56:7 TAG (OR=5.85, 95% CI=2.04-18.02; p=0.001) and C2 carnitine (OR=7.4, 95% CI=2.37-25.35; p=0.001).

The top three metabolites inversely associated with risk were C36:2 phosphatidylcholines (PC) plasmalogen (OR=0.11, 95% CI=0.03-0.35), p=0.0003), C34:1 PC plasmalogen-A (OR=0.18, 95% CI=0.05-0.54, p=0.003), C22:0 lysophosphatidylethanolamine (LPE) (OR=0.21, 95% CI=0.07-0.59; p=0.004). C36:2 PC plasmalogen and pseudouridine had an adjusted p=0.06/p=0.07, respectively. All other metabolites had adjusted p≥0.14. The test of the global null hypothesis that no metabolite was associated with risk had p=0.06.

On the individual metabolite level, histograms and QQ-plots of the nominal p-values (**Supplementary Figure 1)** together with the results of the permutation test suggest the existence of a metabolomic signal for overall ovarian cancer and non-serous tumors.

### Metabolite groups associated with risk of ovarian cancer

In the MSEA analysis, nine metabolite groups were enriched among metabolites associated with risk of ovarian cancer overall at an FDR ≤0.2 (**Figure 2 and Supplementary Table 3**). The top five associated metabolite groups were organic acids and derivatives, PE plasmalogens, TAGs, cholesteryl esters, PC plasmalogens. Nine metabolite groups were associated with risk of serous/PD tumors with FDR ≤0.20 (**Figure 2 and Supplementary Table 3**). The top five associated metabolite groups were: nucleosides, nucleotides and analogues, TAGs, carnitines, sphingomyelins, and alkaloids and derivatives. Finally, eleven metabolite groups were associated with risk of endometrioid/CC tumors at FDR ≤0.20 (**Figure 2 and Supplementary Table 3**). The top five associated metabolite groups were TAGs, DAGs, fatty acyls, lysophosphatidylserines (LPS), and carnitines.

**Figure 2:**
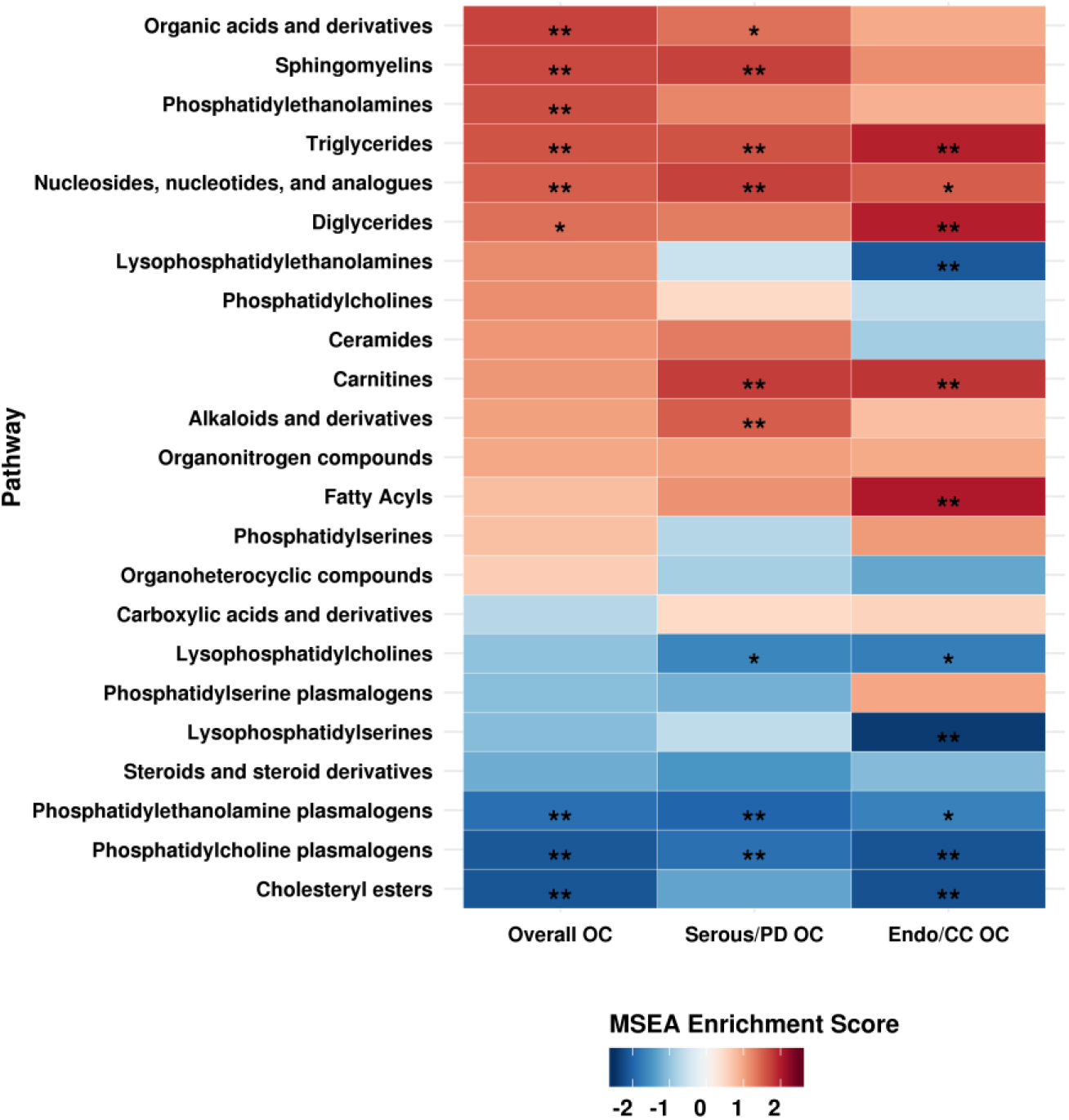
MSEA results. Enriched metabolite groups associated with risk of overall OC, serous/poorly differentiated OC (Serous/PD OC) and endometrioid/clear cell OC (Endo/CC OC). Significance of the association is overlaid on the heat map and marked as follows: * FDR ≤0.2, ** FDR ≤0.05; all other FDR >0.2.

TAGs were enriched among metabolites associated with overall ovarian cancers, serous/PD and endometrioid/CC tumors at FDR≤0.05. Notably, we observed differential associations by acyl carbon number and double bond content with risk of ovarian cancer overall (**Supplementary Figure 2**) and serous/PD tumors (**Supplementary Figure 3**) but not with endometrioid/CC tumors (**Supplementary Figure 4**). Specifically, TAGs with higher number of acyl carbon atoms and double bonds were associated with increased risk, while TAGs with lower number of acyl carbon atoms and double bonds were associated with decreased risk. We did not observe similar patterns for other lipid classes (**Supplementary Figures 2-4**).

### Metabolite modules associated with risk of ovarian cancer

WGCNA identified seven metabolite modules associated with risk of ovarian cancer with FDR ≤0.20 (**Table 3 and Figure 3**). Module 1 (M1, characterized by steroids and steroid derivatives, organic acids and derivatives, and organonitrogen compounds [**Supplementary Figure 5**, **Supplementary Table 4]**), M2 (characterized by TAGs, PCs, PE, LPCs, and LPEs), M6 (characterized by TAGs, LPEs and CEs), and M7 (characterized by TAGs, DAGs, ceramides and CEs) were associated with increased risk of ovarian cancer overall, OR=1.99 (p=0.013/FDR=0.072), 1.62 (p=0.093/FDR=0.186), 1.56 (p=0.081/FDR=0.186) and 1.8 (p=0.015/FDR=0.072), respectively. M4 (characterized by carnitines, pseudouridine [inversely weighted], and organic acids and derivatives) was associated with decreased risk (OR = 0.5, p=0.022/FDR=0.072). M7 (characterized by TAGs, DAGs, ceramides and CEs) was associated with increased risk of serous/PD tumors (OR=1.97; p=0.012/FDR=0.117).

**Table 3:** P-values, FDR, odds ratio (OR) for an increase from the 10^th^ to the 90^th^ percentile of metabolite levels and 95% confidence intervals (CI) of WGCNA metabolite modules associated with risk of ovarian cancer overall and by histotype.

|  | Overall OC |  |  |  | Serous/PD OC |  |  |  | Endometrioid/CC OC |  |  |  |
| --- | --- | --- | --- | --- | --- | --- | --- | --- | --- | --- | --- | --- |
| Module /<br>number<br>of metabolites | OR<br>(95% CI) | P-valu<br>e | FD<br>R | explained<br>variance<br>[%] | OR<br>(95% CI) | P-value | FD<br>R | explained<br>variance<br>[%] | OR<br>(95% CI) | P-value | FDR | explained<br>variance<br>[%] |
| M1 / 24 | 1.99<br>(1.15-3.42) | 0.01<br>3 | 0.07<br>2 | 14.51 | 1.64<br>(0.89-3.05) | 0.114 | 0.46 | 14.87 | 1.09<br>(0.38-3.11) | 0.875 | 0.613 | 14.62 |
| M2 / 79 | 1.62<br>(0.92-2.85) | 0.09<br>3 | 0.18<br>6 | 23.63 | 1.16<br>(0.65-2.07) | 0.624 | 0.70<br>1 | 23.35 | 6.14<br>(1.97-20.67) | 0.002 | 0.011 | 24.54 |
| M3 / 76 | 0.9<br>(0.56-1.46) | 0.67<br>8 | 0.68<br>2 | 65.22 | 1.06<br>(0.62-1.8) | 0.839 | 0.83<br>9 | 65.18 | 0.48<br>(0.17-1.28) | 0.151 | 0.212 | 66.23 |
| M4 / 74 | 0.5<br>(0.28-0.9) | 0.02<br>2 | 0.07<br>2 | 17.86 | 0.64<br>(0.35-1.15) | 0.138 | 0.46 | 18.13 | 0.17<br>(0.05-0.54) | 0.003 | 0.011 | 17.63 |
| M5 / 49 | 0.91<br>(0.56-1.46) | 0.68<br>2 | 0.68<br>2 | 24.23 | 1.2<br>(0.71-2.03) | 0.49 | 0.70<br>1 | 23.94 | 0.35<br>(0.12-0.94) | 0.041 | 0.072 | 21.61 |
| M6 / 37 | 1.56<br>(0.95-2.58) | 0.08<br>1 | 0.18<br>6 | 40.38 | 1.17<br>(0.68-2.01) | 0.575 | 0.70<br>1 | 40.32 | 1.4<br>(0.5-3.99) | 0.522 | 0.456 | 40.16 |
| M7 / 32 | 1.8<br>(1.12-2.88) | 0.01<br>5 | 0.07<br>2 | 32.45 | 1.97<br>(1.17-3.36) | 0.012 | 0.11<br>7 | 32 | 1.94<br>(0.71-5.5) | 0.203 | 0.237 | 32.69 |
| M8 / 24 | 0.82<br>(0.49-1.37) | 0.44<br>1 | 0.55<br>2 | 65.98 | 0.81<br>(0.47-1.4) | 0.451 | 0.70<br>1 | 65.34 | 0.22<br>(0.07-0.65) | 0.007 | 0.017 | 64.15 |
| M9 / 14 | 0.73<br>(0.45-1.19) | 0.20<br>5 | 0.34<br>1 | 37.33 | 0.88<br>(0.51-1.5) | 0.631 | 0.70<br>1 | 38.1 | 0.79<br>(0.29-2.21) | 0.651 | 0.506 | 36.26 |
| M10 / 11 | 0.75<br>(0.43-1.29) | 0.29<br>3 | 0.41<br>9 | 46.53 | 0.7<br>(0.38-1.29) | 0.257 | 0.64<br>4 | 45.92 | 1.88<br>(0.57-6.46) | 0.305 | 0.305 | 47.51 |

**Figure 3:**
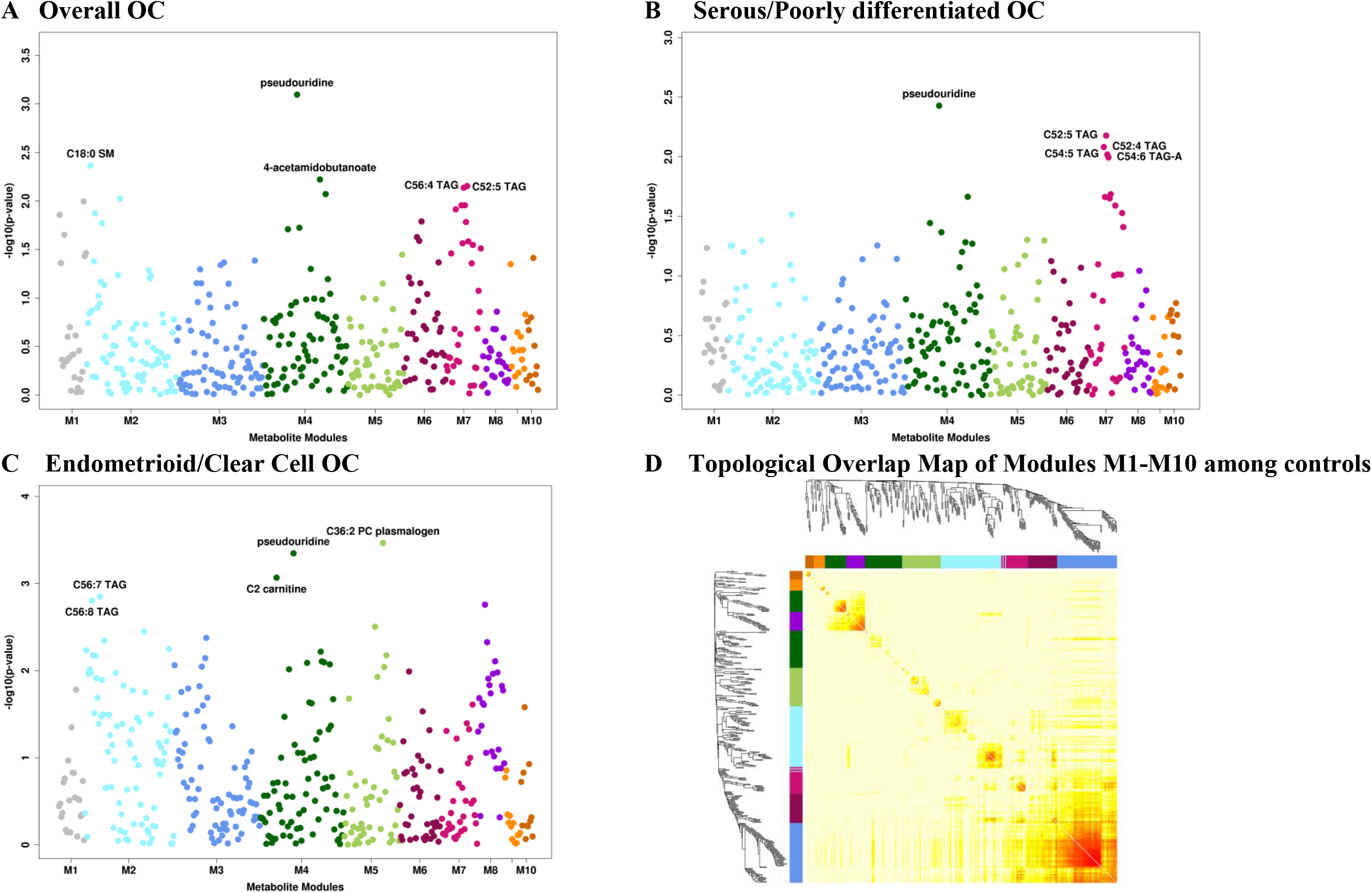
METhattan plots. Man*hattan* plots of *met*abolites by metabolite groups, with each group being shown in a different color. **A** Overall ovarian cancer. **B** Serous/poorly differentiated ovarian cancer. **C** Endometrioid/clear cell ovarian cancer**. D** Topological Overlap Matrix (TOM). Metabolites in the rows and columns are sorted by the clustering tree. Light yellow shades represent low topological overlap (low similarity). Darker red shades represent higher overlap and similarity. Metabolite modules correspond to the squares along the diagonal.

Finally, four modules were associated with risk of endometrioid/CC tumors. M2 was positively associated (OR=6.14; p=0.002/FDR=0.011) with risk while M4 had the strongest inverse association (OR=0.17; p=0.003/FDR=0.011). PC plasmalogens and PE plasmalogens characterized M5 while M8 primarily included fatty acyls, and both were inversely associated with risk.

### Metabolites associated with ovarian cancer risk by menopausal status at blood collection

C22:0 LPS isomer was suggestively associated with increased risk among postmenopausal women (OR=1.83, 95%CI=0.92-3.63; p=0.085) and decreased risk among premenopausal women at blood collection (OR=0.44, 95%CI=0.17-1.08; p=0.074) with a heterogeneity p=0.004 (**Supplementary Table 5**). C38:4 PC plasmalogen was suggestively associated with increased risk among postmenopausal women (OR=1.92, 95%CI=0.96-3.85; p=0.066) and decreased risk among premenopausal women at blood collection (OR=0.16, 95%CI=0.05-0.51; p=0.002) with a heterogeneity p=0.005. 14/22 (63%) metabolites associated with risk (p≤0.1) among premenopausal women but only 15/98 (15%) metabolites associated with risk (p≤0.1) among postmenopausal women showed inverse associations. Pseudouridine did not show heterogeneity by menopausal status (heterogeneity p=0.32).

### Metabolites associated with ovarian cancer risk by time between blood collection and diagnosis

Hydroxyvitamin D3 was associated with increased risk among participants with blood collection 12-23 years after diagnosis (OR=1.84, 95%CI=1.02-3.37; p=0.044) but not among participants with blood collection 3-11 years after diagnosis (OR=0.64, 95%CI=0.36-1.15; p=0.141) with a heterogeneity p=0.002 (**Supplementary Table 6**). C40:6 phosphatidylserine (PS) was associated with decreased risk among participants with blood collection 12-23 years after diagnosis (OR=0.55, 95%CI=0.3-0.99; p=0.049) but not among participants with blood collection 3-11 years after diagnosis (OR=1.41, 95%CI=0.79-2.51; p=0.245) with a heterogeneity p=0.008. Pseudouridine showed suggestively stronger associations (heterogeneity p=0.066) among women for whom sample collection was 3-11 years before diagnosis (OR=4.48, 95%CI=2.25-9.24; p≤0.001) compared to participants with samples collection 12-23 years before diagnosis (OR=2.00, 95%CI=1.06-3.85; p=0.035).

### Metabolites associated with ovarian cancer risk by tumor aggressiveness

Fifty-three lipid-related metabolites (26 TAGs, 7PCs, 6 LPEs, 3 PEs, 3 LPC, 4DAGs, 2 LPSs, and 2 PSs) showed differences by tumor aggressiveness at heterogeneity p≤0.01 (Supplementary Table 7). Seven metabolites (6 TAGs and 1 PSs) were associated with increased risk of rapidly fatal disease with ORs ranging between 2.56 and 3.07 at p≤0.008 but not with risk of less aggressive tumors (p>0.62) with heterogeneity p≤0.001. Several lipid-related metabolite classes (DAGs, LPCs, LPEs, PCs, PEs, PSs, and TAGs with high acyl carbon content and saturation) were upregulated in rapidly fatal tumors while carnitines were up-regulated in less aggressive tumors (Supplementary Figure 8). TAGs with lower acyl carbon content and saturation were inversely associated with less aggressive tumors. Pseudouridine did not show heterogeneity by tumor aggressiveness (heterogeneity p=0.13).

## Discussion

We conducted the first large-scale agnostic analysis of metabolomics and risk of ovarian cancer. We identified a potential novel risk factor, plasma pseudouridine, which was associated with an increased risk of ovarian cancer overall and non-serous tumors. Stronger associations for pseudouridine were observed among cases diagnosed within 3-11 years after blood collection. We identified several metabolite groups and metabolite modules associated with risk of ovarian cancer risk, as well as multiple subtype-specific associations, that open up new opportunities for assessing novel metabolite pathways involved in ovarian cancer risk.

### Pseudouridine

Pseudouridine is a post-transcriptionally modified nucleoside, and is an isomer of uridine. Cellular RNA contains more than 100 modified nucleosides with pseudouridine being the most abundant (23). Pseudouridine is produced by pseudouridine synthase by isomerizing uridines from transfer RNA (24), which is involved in in protein translation, and from spliceosomal snRNA which plays a role in pre-mRNA splicing (25). Pseudouridine was associated with risk of ovarian cancer and non-serous tumors but not serous tumors. The magnitude of the association between pseudouridine and serous cancer was similar to that of overall ovarian cancer risk though not significant, while the risk estimate for endometrioid/CC tumors was higher. This is likely due, in part, to limited sample sizes in the histotype-specific analyses. Our results suggest that pseudouridine may represent a common etiologic mechanism underlying different histotypes of ovarian cancer, which has been observed for other risk factors, such as aspirin and CRP (11, 26, 27). In retrospective studies, pseudouridine was elevated in urine (28) and plasma (29) from epithelial ovarian cancer patients compared to healthy controls. This, in combination with our finding that pseudouridine had a stronger association when assessed 3-11 years before diagnosis, suggests that this modified nucleotide may be important in progression of preclinical lesions to fully overt invasive disease, which for high-grade serous ovarian cancer appears to be about 7-9 years (30, 31).

Increasing evidence suggests that pseudouridylation plays a role in cancer-associated splicing distributions (25, 32), which are more variable than in normal tissues, and in which tissue-specific alternative splicing reverts to a default cancer pattern that directly contributes to cellular transformation and cancer progression (33, 34). This has been observed in serous carcinomas, which have highly dysregulated splicing compared to normal tissue (35). Further, aberrant pseudouridylation may lead to altered translation and reduced translational fidelity of p53 (36, 37), which is mutated in nearly all high-grade serous tumors (38). Another potential mechanism is via circular RNA activity, which is altered due to isomerization of uridine to pseudouridine, and has been shown to be dysregulated in ovarian cancer (39). Additional research should explore the potential role of pseudouridine in precursor lesions to ovarian cancer and the relation between circulating pseudouridine to ovarian and fallopian tube tissue levels.

### Triacylglycerides

Notably, several individual TAGs were nominally related to risk and showed significant heterogeneity by tumor aggressiveness (increased circulating TAG levels were associated with increased risk of rapidly fatal tumors but not less aggressive tumors). TAGs as a group were enriched in the MSEA analysis, and 3 of 7 WCGNA modules related to risk were characterized by TAGs. Long chain fatty acids, a main source of energy in the human body, are stored and transported from the small intestine and liver to peripheral cells in the form of TAGs (40). Lipid synthesis and metabolism, specifically release of free fatty acids from TAGs, are dysregulated in ovarian tumors, increasing cell migration and invasive potential (41–44). Further, several human studies reported suggestive associations of ovarian cancer risk with total cholesterol (45) (positive) or HDL (inverse) (46). Additionally, evidence has demonstrated that ovarian cancer metastasizes preferentially to the adipose-rich omentum, a key player in the creation of the metastatic tumor microenvironment in the intraperitoneal cavity (47). Omental fat possess a distinct lipidomic signature with several lipid groups, including TAGs, DAGs, and SMs, showing differences when compared to subcutaneous fat (48). Finally, plasma TAGs represent known risk factors for cardiovascular disease (49) and coronary heart disease (40). A recent study identified that TAGs at the extremes of carbon atoms and saturation had differential associations with diabetes risk (50). We also observed differential associations by TAG fatty acids length and saturation, with higher number of carbon atoms and double bonds related to an increased risk and lower number of carbon atoms and double bonds related to decreased risk, particularly for serous/PD tumors. A similar pattern was observed in a retrospective study of serum samples from high-grade serous ovarian cancer cases and controls (51). Together with our results, these findings suggest that circulating TAG levels may be a risk biomarker for ovarian cancer, particularly for rapidly fatal tumors. Additional prospective studies are needed to validate these associations in different populations and assess the potential differential role of various TAG species in ovarian carcinogenesis.

### Other metabolite groups

A number of metabolite groups and clusters were associated with ovarian cancer risk, including organic acids and derivatives, and SMs, the latter of which was hypothesized a priori as a potential risk biomarker and is discussed elsewhere (Zeleznik el al., in revision (52)). A metabolite module driven by carnitines, organic acids and derivatives, carboxylic acids and derivatives, which included pseudouridine (highly negatively weighted) was associated with decreased risk of overall ovarian cancer and non-serous tumors. This module includes asymmetric dimethylarginine (ADMA), which has been related to risk of cardiovascular disease (53), and inhibits nitric oxide synthesis and may have antiproliferative properties (54); nitric oxide signaling may be involved in ovarian carcinogenesis (55). LPEs were also represented in two WCGNA modules associated with increased risk of overall ovarian cancer, and one WCGNA module associated with increased risk of endometrioid and CC tumors. LPEs have been shown to increase migration in response to chemotherapy as well as have invasive potential in ovarian cancer cell lines (56). In MSEA analyses, several metabolite classes had a significant inverse enrichment score, including PE plasmalogens, PC plasmalogens and cholesteryl esters, independent of subtype. PE plasmalogens and PC plasmalogens were highly weighted in the WGCNA-derived module M5 which was inversely associated with risk of endometrioid/CC tumors while cholesteryl esters where highly negatively weighted in M7 which was associated with increased risk of overall ovarian cancer and serous/PD tumors. Notably, C36:2 PC plasmalogen was associated with lower risk of endometrioid/CC tumors (OR=0.11, 95%CI =0.03-0.35; permutation adjusted p=0.056). Little work has examined these markers in ovarian cancer development or etiology.

Our study has several strengths and limitations. Importantly, this is a prospective metabolomics study of ovarian cancer risk with coverage of multiple different metabolite classes. While we had over 250 total ovarian cancer cases and controls, we had more limited sample sizes for specific histotypes, which have been shown to have different associations for known risk factors (57). To maximize our power, borderline and tumors of unknown morphology were analyzed together with invasive tumors. We did not include information on family history of ovarian cancer. However, only 2 of the 252 cases were diagnosed before age 45 suggesting that early onset disease, likely due to high risk mutations, does not play a role in this study. We also applied stringent QC criteria to limit identification of spurious associations. Additional strengths include the long follow-up time and detailed covariate information. A limitation is that we only analyzed blood samples collected at one point in time; however, we previously showed that the majority of the measured metabolites have a high within person stability over time (15). Further, we do not have an independent validation dataset. As this type of data becomes more common, further population studies are needed to validate the results discussed here, while experimental studies are required to understand the biological mechanisms underlying these associations.

In summary, circulating levels of plasma pseudouridine were associated with higher risk of ovarian cancer 3-23 years before diagnosis, with stronger associations among participants with samples collected closer to diagnosis. Additionally, several metabolite groups and metabolite modules were associated with risk of disease, independent of subtype as well as subtype specific. While independent prospective studies are needed for validation, our results highlight some potentially important novel metabolites that may play a role in the etiology of ovarian cancer. Further work is warranted to explore the potential use of these metabolites as targets for prevention and/or predictors of risk of ovarian cancer.

## Acknowledgements

This study was funded by the National Cancer Institute (P01 CA087969, U01 CA176726 UM1 CA186107) and the Department of Defense (W81XWH-12-1-0561). We would like to thank the participants and staff of the Nurses Health Studies and Nurses Health Studies II for their valuable contributions as well as the following state cancer registries for their help: AL, AZ, AR, CA, CO, CT, DE, FL, GA, ID, IL, IN, IA, KY, LA, ME, MD, MA, MI, NE, NH, NJ, NY, NC, ND, OH, OK, OR, PA, RI, SC, TN, TX, VA, WA, WY. The authors assume full responsibility for analyses and interpretation of these data.

## Supplementary Figures

**Supplementary Figure 1:**
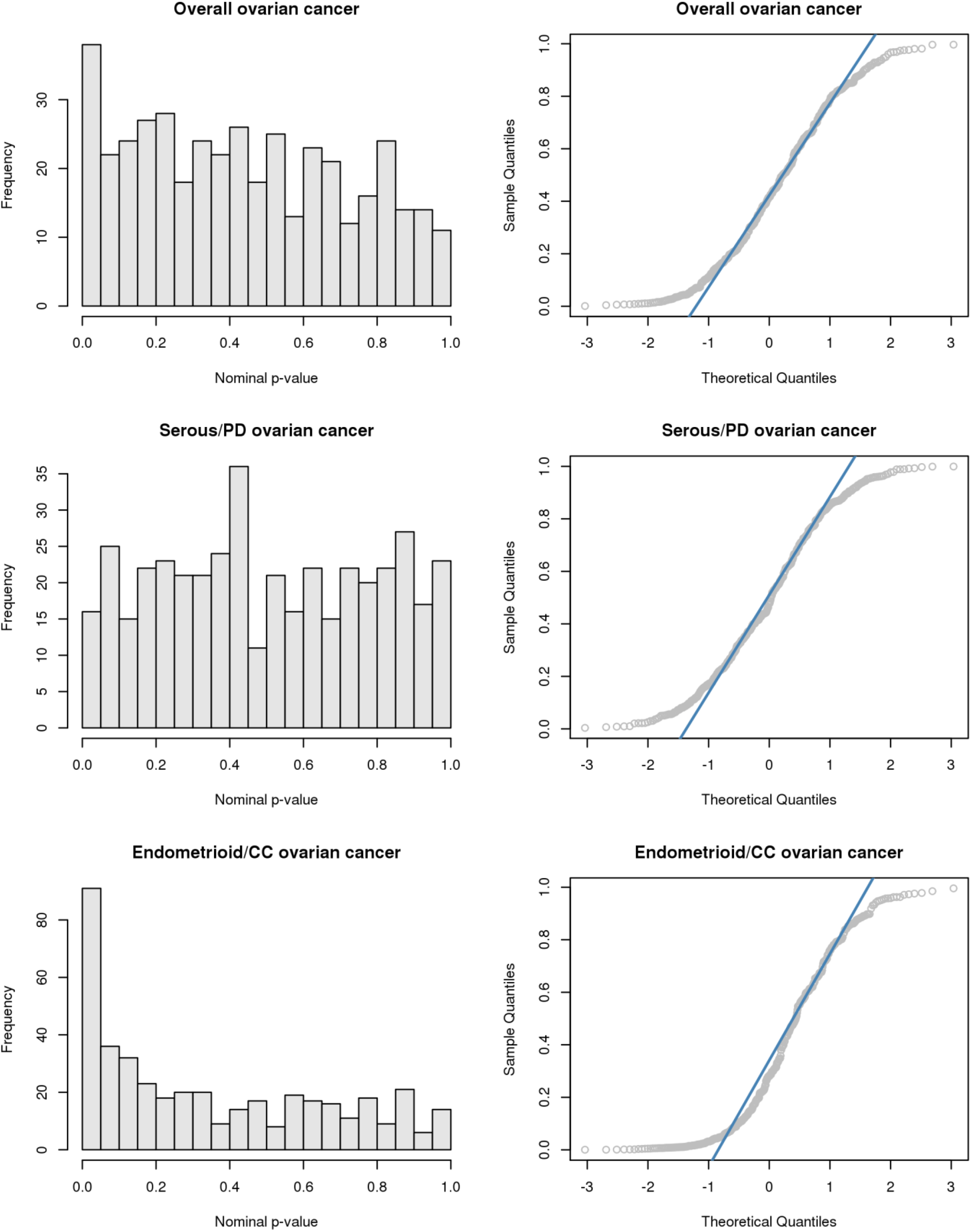
Histograms and QQ-plots of the nominal p-values for overall ovarian cancer and by histotype.

**Supplementary Figure 2:**
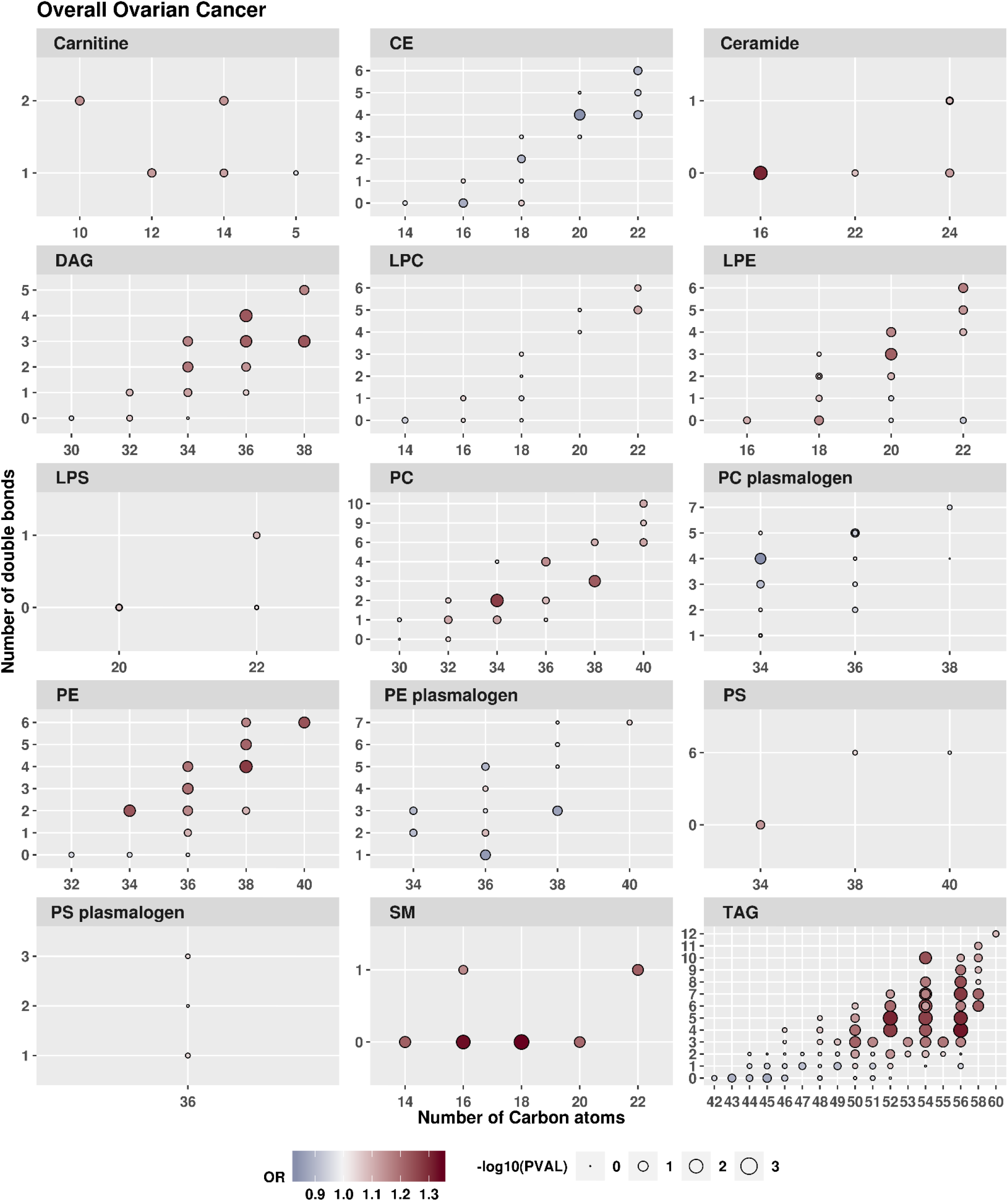
Odds ratio (OR) of overall ovarian cancer for an increase from the 10^th^ to the 90^th^ percentile of metabolite levels. Results are shown by metabolite group, by the number of Carbon atoms and by the number of double bonds.

**Supplementary Figure 3:**
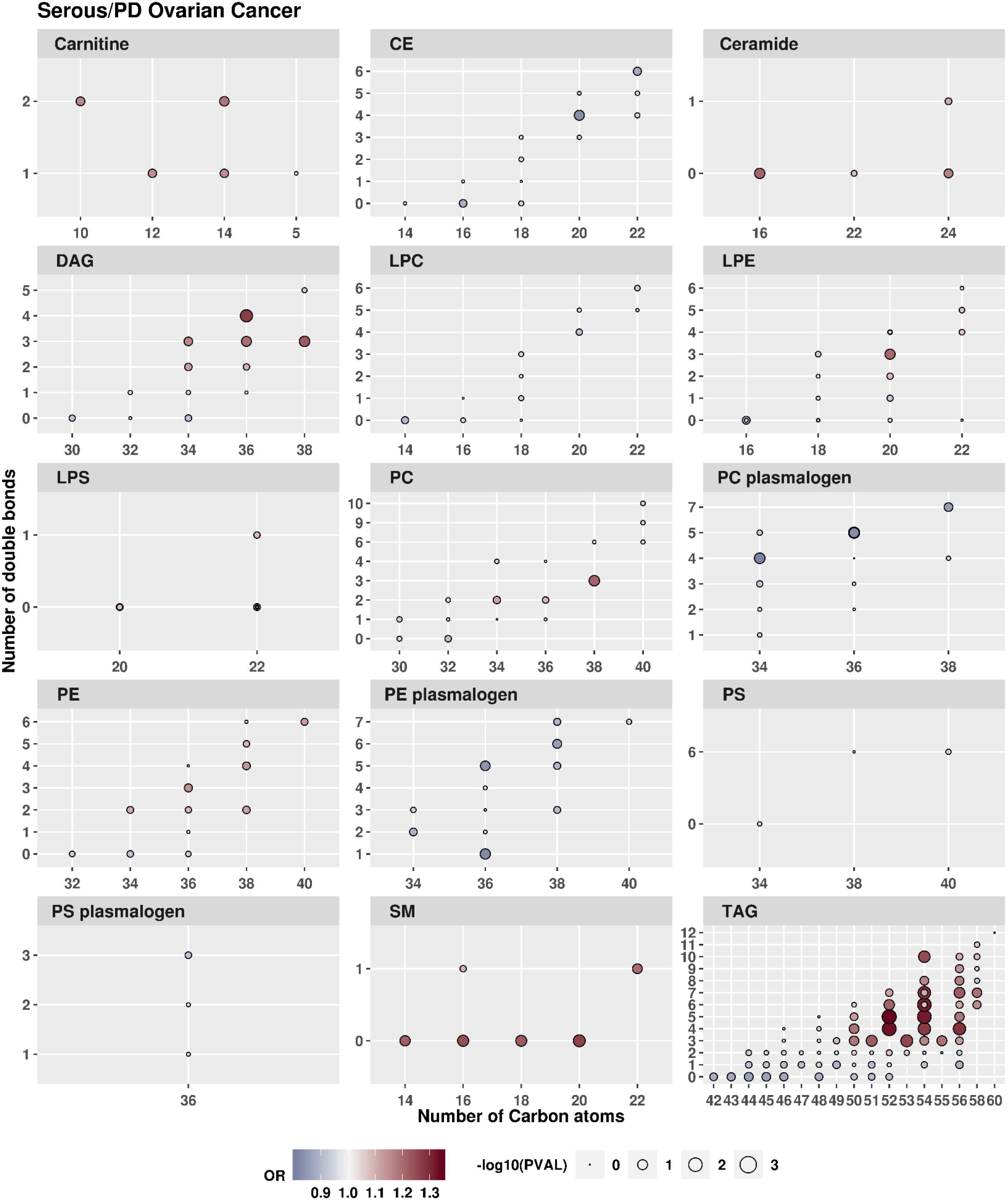
Odds ratio (OR) of serous/PD ovarian tumors for an increase from the 10^th^ to the 90^th^ percentile of metabolite levels. Results are shown by metabolite group, by the number of Carbon atoms and by the number of double bounds.

**Supplementary Figure 4:**
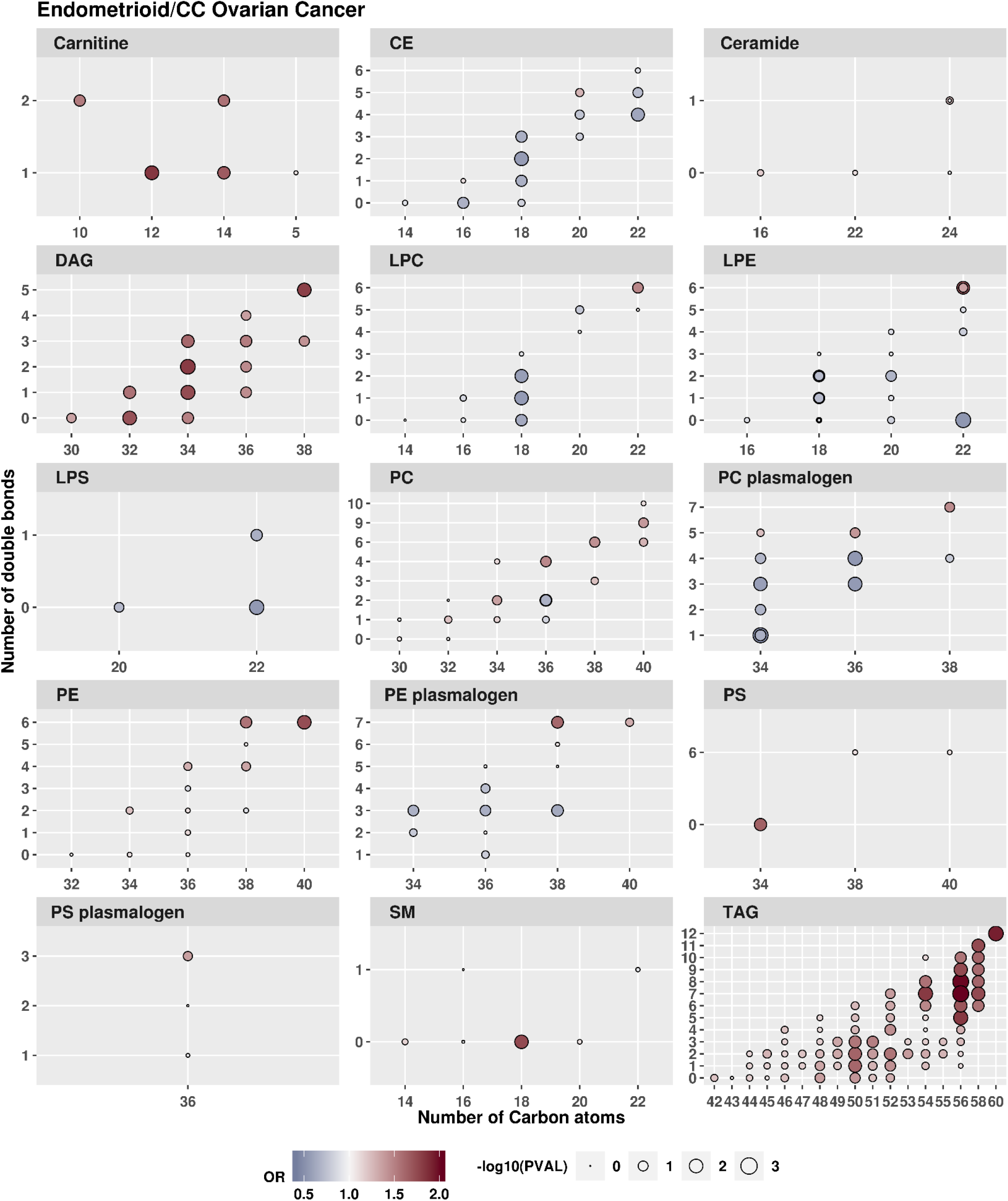
Odds ratio (OR) of endometrioid/CC ovarian tumors for an increase from the 10^th^ to the 90^th^ percentile of metabolite levels. Results are shown by metabolite group, by the number of Carbon atoms and by the number of double bounds.

**Supplementary Figure 5:**
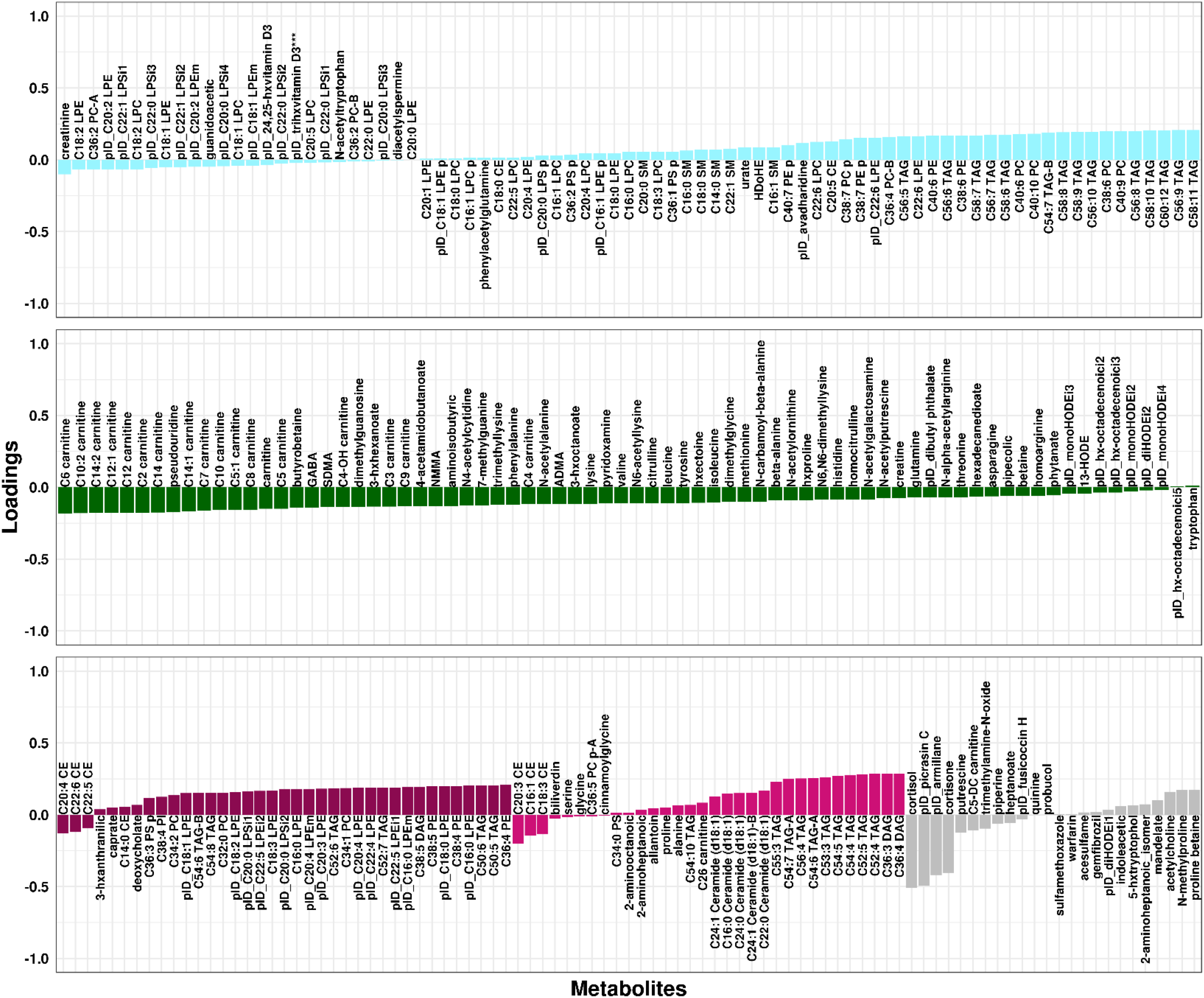
WGCNA module loadings for overall ovarian cancer. Shown are loadings from modules associated with overall ovarian cancer at FDR<0.2. Loadings/module colors correspond to colors in Figure 3.

**Supplementary Figure 6:**
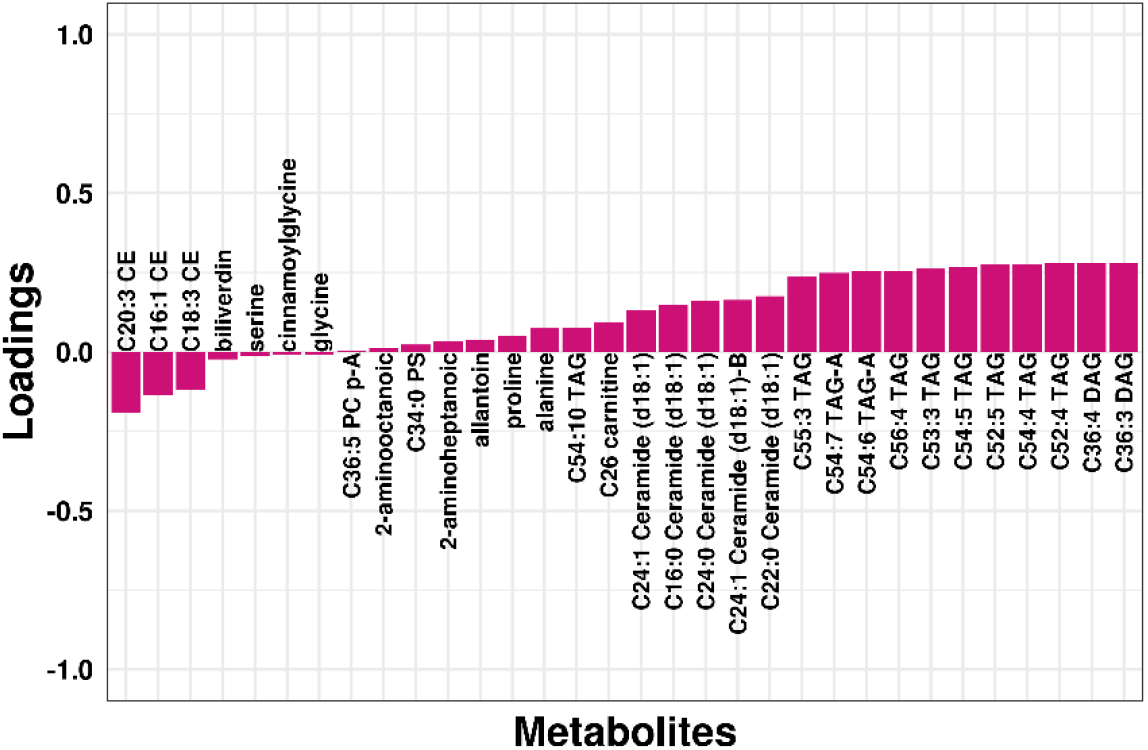
WGCNA module loadings for serous/PD ovarian cancer. Shown are loadings from modules associated with serous/PD tumors at FDR<0.2. Loadings/module colors correspond to colors in Figure 3.

**Supplementary Figure 7:**
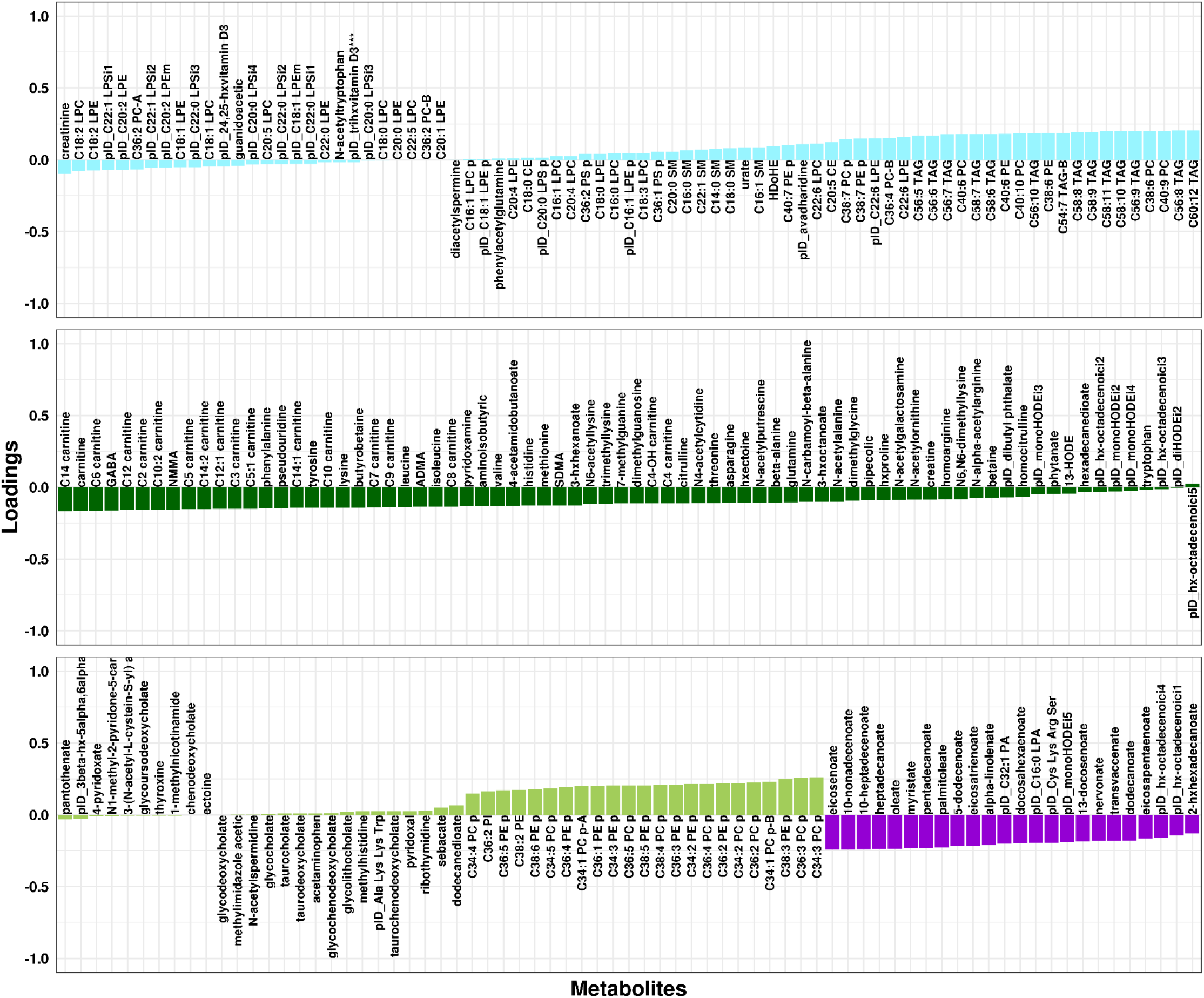
WGCNA module loadings for endometrioid/CC ovarian cancer. Shown are loadings from modules associated with endometrioid/CC tumors at FDR<0.2. Loadings/module colors correspond to colors in Figure 3.

**Supplementary Figure 8:**
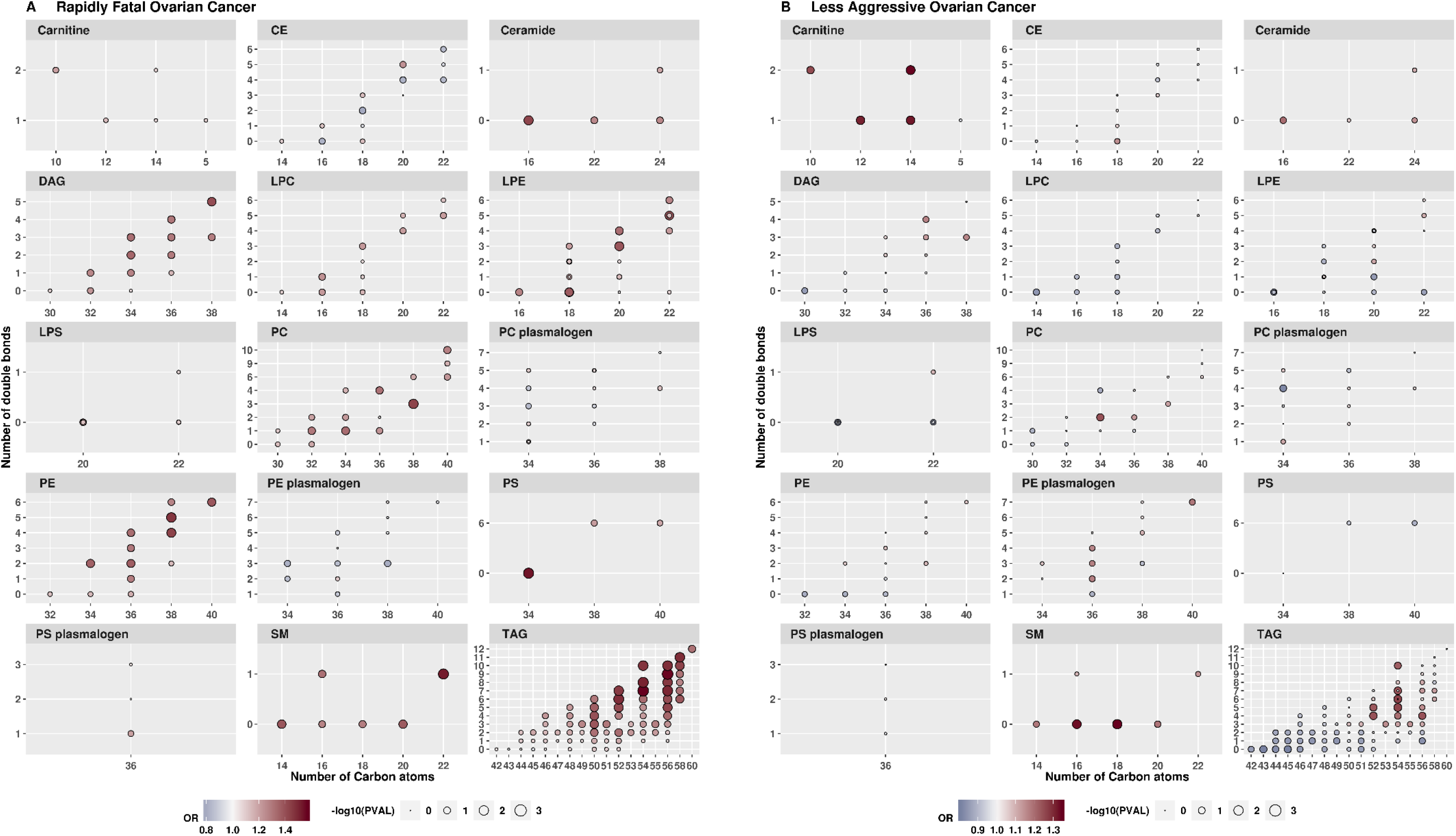
Odds ratio (OR) of rapidly fatal and less aggressive ovarian tumors for an increase from the 10^th^ to the 90^th^ percentile of metabolite levels. Results are shown by metabolite group, by the number of Carbon atoms and by the number of double bounds.

## References

1. American Cancer Society. Cancer Facts & Figures 2018. Atlanta: American Cancer Society; 2018.

2. Krumsiek J, Bartel J, Theis FJ. Computational approaches for systems metabolomics. Curr Opin Biotechnol. 2016;39:198–206.

3. Goodacre R. Making sense of the metabolome using evolutionary computation: seeing the wood with the trees. J Exp Bot. 2005;56(410):245–54.

4. Shah SH, Kraus WE, Newgard CB. Metabolomic profiling for the identification of novel biomarkers and mechanisms related to common cardiovascular diseases: form and function. Circulation. 2012;126(9):1110–20.

5. Mayers JR, Wu C, Clish CB, Kraft P, Torrence ME, Fiske BP, et al. Elevation of circulating branched-chain amino acids is an early event in human pancreatic adenocarcinoma development. Nat Med. 2014;20(10):1193.

6. Mondul AM, Moore SC, Weinstein SJ, Karoly ED, Sampson JN, Albanes D. Metabolomic analysis of prostate cancer risk in a prospective cohort: The alpha‐tocopherol, beta‐carotene cancer prevention (ATBC) study. Int J Cancer. 2015;137(9):2124–32.

7. Playdon MC, Ziegler RG, Sampson JN, Stolzenberg-Solomon R, Thompson HJ, Irwin ML, et al. Nutritional metabolomics and breast cancer risk in a prospective study. The American journal of clinical nutrition. 2017;106(2):637–49.

8. Moore SC, Playdon MC, Sampson JN, Hoover RN, Trabert B, Matthews CE, et al. A metabolomics analysis of body mass index and postmenopausal breast cancer risk. JNCI: Journal of the National Cancer Institute. 2018.

9. Hankinson SE, Willett WC, Michaud DS, Manson JE, Colditz GA, Longcope C, et al. Plasma prolactin levels and subsequent risk of breast cancer in postmenopausal women. J Natl Cancer Inst. 1999;91(7):629–34.

10. Tworoger SS, Sluss P, Hankinson SE. Association between plasma prolactin concentrations and risk of breast cancer among predominately premenopausal women. Cancer Res. 2006;66(4):2476–82.

11. Poole EM, Lee I-M, Ridker PM, Buring JE, Hankinson SE, Tworoger SS. A prospective study of circulating C-reactive protein, interleukin-6, and tumor necrosis factor α receptor 2 levels and risk of ovarian cancer. American journal of epidemiology. 2013;178(8):1256–64.

12. Mascanfroni ID, Takenaka MC, Yeste A, Patel B, Wu Y, Kenison JE, et al. Metabolic control of type 1 regulatory T cell differentiation by AHR and HIF1-α. Nat Med. 2015;21(6):638.

13. O’sullivan JF, Morningstar JE, Yang Q, Zheng B, Gao Y, Jeanfavre S, et al. Dimethylguanidino valeric acid is a marker of liver fat and predicts diabetes. The Journal of clinical investigation. 2017;127(12):4394–402.

14. Paynter NP, Balasubramanian R, Giulianini F, Wang DD, Tinker LF, Gopal S, et al. Metabolic predictors of incident coronary heart disease in women. Circulation. 2018;137(8):841–53.

15. Townsend MK, Clish CB, Kraft P, Wu C, Souza AL, Deik AA, et al. Reproducibility of metabolomic profiles among men and women in 2 large cohort studies. Clinical chemistry. 2013;59(11):1657–67.

16. Westfall PH, Young SS. Resampling-based multiple testing: Examples and methods for p-value adjustment: John Wiley & Sons; 1993.

17. Subramanian A, Tamayo P, Mootha VK, Mukherjee S, Ebert BL, Gillette MA, et al. Gene set enrichment analysis: a knowledge-based approach for interpreting genome-wide expression profiles. Proceedings of the National Academy of Sciences. 2005;102(43):15545–50.

18. Sergushichev A. An algorithm for fast preranked gene set enrichment analysis using cumulative statistic calculation. BioRxiv. 2016:060012.

19. Langfelder P, Horvath S. WGCNA: an R package for weighted correlation network analysis. BMC Bioinformatics. 2008;9(1):559.

20. Barabasi A-L, Oltvai ZN. Network biology: understanding the cell’s functional organization. Nature reviews genetics. 2004;5(2):101–13.

21. Benjamini Y, Krieger AM, Yekutieli D. Adaptive linear step-up procedures that control the false discovery rate. Biometrika. 2006;93(3):491–507.

22. Team RC. R: A language and environment for statistical computing. R Foundation for Statistical Computing, Vienna, Austria. 2013. 2014.

23. Hamma T, Ferré-D’Amaré AR. Pseudouridine synthases. Chem Biol. 2006;13(11):1125–35.

24. Johnson L, Söll D. In vitro biosynthesis of pseudouridine at the polynucleotide level by an enzyme extract from Escherichia coli. Proceedings of the National Academy of Sciences. 1970;67(2):943–50.

25. Yu Y-T, Meier UT. RNA-guided isomerization of uridine to pseudouridine— pseudouridylation. RNA Biol. 2014;11(12):1483–94.

26. Ose J, Schock H, Tjønneland A, Hansen L, Overvad K, Dossus L, et al. Inflammatory markers and risk of epithelial ovarian cancer by tumor subtypes: the EPIC cohort. Cancer Epidemiology and Prevention Biomarkers. 2015;24(6):951–61.

27. Trabert B, Ness RB, Lo-Ciganic W-H, Murphy MA, Goode EL, Poole EM, et al. Aspirin, nonaspirin nonsteroidal anti-inflammatory drug, and acetaminophen use and risk of invasive epithelial ovarian cancer: a pooled analysis in the Ovarian Cancer Association Consortium. JNCI: Journal of the National Cancer Institute. 2014;106(2).

28. Zhang T, Wu X, Ke C, Yin M, Li Z, Fan L, et al. Identification of potential biomarkers for ovarian cancer by urinary metabolomic profiling. J Proteome Res. 2012;12(1):505–12.

29. Ke C, Hou Y, Zhang H, Fan L, Ge T, Guo B, et al. Large‐scale profiling of metabolic dysregulation in ovarian cancer. Int J Cancer. 2015;136(3):516–26.

30. Conner JR, Meserve E, Pizer E, Garber J, Roh M, Urban N, et al. Outcome of unexpected adnexal neoplasia discovered during risk reduction salpingo-oophorectomy in women with germ-line BRCA1 or BRCA2 mutations. Gynecol Oncol. 2014;132(2):280–6.

31. Labidi-Galy SI, Papp E, Hallberg D, Niknafs N, Adleff V, Noe M, et al. High grade serous ovarian carcinomas originate in the fallopian tube. Nature communications. 2017;8(1):1093.

32. Zhao Y, Dunker W, Yu Y-T, Karijolich J. The Role of Noncoding RNA Pseudouridylation in Nuclear Gene Expression Events. Frontiers in Bioengineering and Biotechnology. 2018;6:8.

33. Venables JP, Klinck R, Koh C, Gervais-Bird J, Bramard A, Inkel L, et al. Cancer-associated regulation of alternative splicing. Nature structural & molecular biology. 2009;16(6):670.

34. El Marabti E, Younis I. The Cancer Spliceome: reprograming of alternative splicing in cancer. Frontiers in molecular biosciences. 2018;5.

35. Klinck R, Bramard A, Inkel L, Dufresne-Martin G, Gervais-Bird J, Madden R, et al. Multiple alternative splicing markers for ovarian cancer. Cancer research. 2008;68(3):657–63.

36. Ji B, Harris B, Liu Y, Deng Y, Gradilone S, Cleary M, et al. Targeting IRES-mediated p53 synthesis for cancer diagnosis and therapeutics. International journal of molecular sciences. 2017;18(1):93.

37. Penzo M, Guerrieri A, Zacchini F, Treré D, Montanaro L. RNA pseudouridylation in physiology and medicine: for better and for worse. Genes. 2017;8(11):301.

38. Kanchi KL, Johnson KJ, Lu C, McLellan MD, Leiserson MD, Wendl MC, et al. Integrated analysis of germline and somatic variants in ovarian cancer. Nature communications. 2014;5:3156.

39. Kristensen L, Hansen T, Venø M, Kjems J. Circular RNAs in cancer: opportunities and challenges in the field. Oncogene. 2018;37(5):555.

40. Austin MA. Plasma triglyceride as a risk factor for coronary heart disease. The epidemiologic evidence and beyond. Am J Epidemiol. 1989;129(2):249–59.

41. Pyragius CE, Fuller M, Ricciardelli C, Oehler MK. Aberrant lipid metabolism: an emerging diagnostic and therapeutic target in ovarian cancer. Int J Mol Sci. 2013;14(4):7742–56.

42. Tania M, Khan MA, Song Y. Association of lipid metabolism with ovarian cancer. Curr Oncol. 2010;17(5):6–11.

43. Nomura DK, Long JZ, Niessen S, Hoover HS, Ng S-W, Cravatt BF. Monoacylglycerol lipase regulates a fatty acid network that promotes cancer pathogenesis. Cell. 2010;140(1):49–61.

44. Chakraborty PK, Xiong X, Mustafi SB, Saha S, Dhanasekaran D, Mandal NA, et al. Role of cystathionine beta synthase in lipid metabolism in ovarian cancer. Oncotarget. 2015;6(35):37367.

45. Helzlsouer KJ, Alberg AJ, Norkus EP, Morris JS, Hoffman SC, Comstock GW. Prospective study of serum micronutrients and ovarian cancer. J Natl Cancer Inst. 1996;88(1):32–7.

46. Melvin JC, Seth D, Holmberg L, Garmo H, Hammar N, Jungner I, et al. Lipid profiles and risk of breast and ovarian cancer in the Swedish AMORIS study. Cancer Epidemiol Biomarkers Prev. 2012;21(8):1381–4.

47. Motohara T, Masuda K, Morotti M, Zheng Y, El-Sahhar S, Chong KY, et al. An evolving story of the metastatic voyage of ovarian cancer cells: cellular and molecular orchestration of the adipose-rich metastatic microenvironment. Oncogene. 2018:1.

48. Jové M, Moreno-Navarrete JM, Pamplona R, Ricart W, Portero-Otín M, Fernández-Real JM. Human omental and subcutaneous adipose tissue exhibit specific lipidomic signatures. The FASEB Journal. 2014;28(3):1071–81.

49. Hokanson JE, Austin MA. Plasma triglyceride level is a risk factor for cardiovascular disease independent of high-density lipoprotein cholesterol level: a metaanalysis of population-based prospective studies. J Cardiovasc Risk. 1996;3(2):213–9.

50. Al-Sulaiti H, Diboun I, Banu S, Al-Emadi M, Amani P, Harvey TM, et al. Triglyceride profiling in adipose tissues from obese insulin sensitive, insulin resistant and type 2 diabetes mellitus individuals. J Transl Med. 2018;16(1):175.

51. Braicu EI, Darb-Esfahani S, Schmitt WD, Koistinen KM, Heiskanen L, Pöhö P, et al. High-grade ovarian serous carcinoma patients exhibit profound alterations in lipid metabolism. Oncotarget. 2017;8(61):102912.

52. Zeleznik OA, Clish CB, Kraft P, Avila-Pancheco J, Eliassen AH, Tworoger SS. Circulating Lysophosphatidylcholines, Phosphatidylcholines, Ceramides, and Sphingomyelins and Ovarian Cancer Risk: a 23-year Prospective Study. bioRxiv. 2019:565044.

53. Krzyzanowska K, Mittermayer F, Wolzt M, Schernthaner G. ADMA, cardiovascular disease and diabetes. diabetes research and clinical practice. 2008;82:S122–S6.

54. Lüneburg N, Harbaum L, Hennigs JK. The endothelial ADMA/NO pathway in hypoxia-related chronic respiratory diseases. BioMed research international. 2014;2014.

55. El-Sehemy A, Postovit L-M, Fu Y. Nitric oxide signaling in human ovarian cancer: a potential therapeutic target. Nitric Oxide. 2016;54:30–7.

56. Park KS, Lee HY, Lee SY, Kim M-K, Kim SD, Kim JM, et al. Lysophosphatidylethanolamine stimulates chemotactic migration and cellular invasion in SK‐ OV3 human ovarian cancer cells: Involvement of pertussis toxin‐sensitive G‐protein coupled receptor. FEBS letters. 2007;581(23):4411–6.

57. Wentzensen N, Poole EM, Trabert B, White E, Arslan AA, Patel AV, et al. Ovarian cancer risk factors by histologic subtype: an analysis from the Ovarian Cancer Cohort Consortium. J Clin Oncol. 2016;34(24):2888–98.

